# Activity-based CRISPR Scanning Uncovers Allostery in DNA Methylation Maintenance Machinery

**DOI:** 10.1101/2022.05.14.491958

**Authors:** Kevin C. Ngan, Samuel M. Hoenig, Pallavi M. Gosavi, David A. Tanner, Nicholas Z. Lue, Emma M. Garcia, Ceejay Lee, Brian B. Liau

## Abstract

Allostery enables dynamic control of protein function. A paradigmatic example is the tightly orchestrated process of DNA methylation maintenance. Despite their fundamental importance, systematic identification of allosteric sites remains highly challenging. Here we perform CRISPR scanning on the essential maintenance methylation machinery—DNMT1 and its partner UHRF1—with the activity-based inhibitor decitabine to uncover allosteric mechanisms regulating DNMT1. Through computational analyses, we identify putative mutational hotspots in DNMT1 distal from the active site that encompass mutations spanning a multi-domain autoinhibitory interface and the uncharacterized BAH2 domain. We biochemically characterize these mutations as gain-of-function mutations that increase DNMT1 activity. Extrapolating our analysis to UHRF1, we discern putative gain-of-function mutations in multiple domains, including key residues across the autoinhibitory TTD–PBR interface. Collectively, our findings highlight the utility of activity-based CRISPR scanning for nominating candidate allosteric sites, even beyond the direct drug target.

## Introduction

Allostery is a fundamental property that enables proteins to translate the effect of a stimulus at one site to modulate function at another distal site. Despite intense study, the identification of allosteric sites across diverse protein targets remains challenging and highly contextual. Unlike orthosteric sites, allosteric sites are often less conserved between related proteins and the principles governing their structural features and properties are not as well understood.^1,2^ Due to these challenges, fewer experimental and computational approaches exist to identify and characterize allosteric sites.^3^ Nevertheless, there are significant efforts to develop small-molecule allosteric modulators as the structural diversity of allosteric sites offer the potential for greater selectivity, lower toxicity, and fine-tuning of protein function compared to orthosteric ligands.^1,2^ Therefore, the development of new tools that enable the identification of allosteric mechanisms would further our understanding of protein regulation and facilitate drug discovery.

Leveraging pharmacological and genetic perturbations in tandem has been widely successful for target deconvolution and elucidating drug mechanism of action.^4^ In particular, the identification of drug resistance-conferring mutations provides critical validation of on-target engagement and can often illuminate underlying biology.^5^ Although many resistance mutations occur proximally to the drug binding site, they can also arise at distal positions of a target protein. These distal mutations can operate by perturbing allosteric mechanisms, even if the drug binds within the orthosteric site.^6–8^ For example, resistance mutations to ABL1 inhibitors, including both orthosteric and allosteric inhibitors, consistently arise outside the drug binding site and drive resistance by destabilizing the inactive conformation or otherwise neutralizing ABL1 autoinhibition.^8–12^ Such findings raise the prospect that identifying distal drug resistance mutations, either in the direct target or in interacting partners, can be exploited to systematically discover and characterize allosteric mechanisms.

Recently, we and others have used CRISPR–Cas9 tiling mutagenesis screens to uncover modes of small-molecule action by identifying drug resistance mutations.^13–15^ In our approach, termed CRISPR-suppressor scanning, Cas9 is used to systematically mutate a target protein with a pooled library of single-guide RNAs (sgRNAs) spanning its entire coding sequence (CDS) to generate large numbers of diverse protein variants in situ. This surviving cellular pool is then treated with small-molecule inhibitors to identify variants conferring drug resistance. Because resistance mutations can occur beyond the drug binding site, we posited that such mutations might operate by disrupting interactions involved in allosteric regulation of protein function.^14,15^ Consequently, we considered whether the use of an activity-based inhibitor that closely resembles the target protein’s native substrate could be exploited to explicitly identify distal resistance mutations and potential allosteric mechanisms.

## Results

### Activity-based CRISPR scanning nominates distal allosteric sites of DNMT1 and UHRF1

Novel approaches to study allostery are particularly attractive for chromatin-modifying enzymes, whose activities are strictly regulated for proper gene expression. A paradigmatic example is the establishment and maintenance of DNA methylation by DNA methyltransferases (DNMTs). Canonically, DNMT1 performs maintenance methylation, ensuring faithful propagation of the methylation landscape.^16^ Accordingly, DNMT1 activity is tightly controlled to prevent aberrant methylation.^17–21^ While structural and biochemical studies have uncovered important mechanistic details underlying DNMT1 regulation, in vitro studies cannot recapitulate the complexities of the cellular environment and dependence on cofactors such as UHRF1 for DNA methylation in vivo.^22– 26^ These discrepancies highlight outstanding gaps in our understanding of DNMT1 regulation and underscore the importance of investigating allosteric regulatory mechanisms within their endogenous context.

We reasoned that activity-based CRISPR scanning of *DNMT1* and *UHRF1* with decitabine (DAC) could uncover mechanisms of DNMT1 allosteric regulation. DAC is a clinically approved DNA hypomethylating agent that acts through mechanism-based inhibition of DNMTs.^27,28^ DAC is a near-identical analog of cytidine—differing only by two atoms—that is incorporated into genomic DNA during replication. When DNMT1 methylates DAC on the nascent strand, DAC’s unique structure prevents DNMT1 release, forming covalent DNMT1–DNA adducts that drive DNMT1 degradation and subsequent global DNA hypomethylation (**Fig. 1a, Supplementary Fig. 1a,b**). At higher DAC doses, these adducts cause acute cytotoxicity independent of hypomethylation.^29^ Consistent with this mechanism, previous work has shown that reducing DNMT1 protein levels alleviates DAC-induced cytotoxicity by decreasing DNMT1–DNA crosslinks.^30^ Because maintenance methylation is an essential process, we reasoned that mutations in DNMT1 arising in response to DAC treatment would be subject to the following constraints: (1) active site mutations disrupting DAC binding but not substrate recognition are highly unlikely due to its near-identical resemblance to DNMT1’s native substrate, cytidine; (2) loss-of-function mutations alleviating adduct-induced cytotoxicity by reducing *DNMT1* copy number may incur fitness penalties from defects in maintenance methylation. Consequently, such loss-of-function mutations may accompany hypermorphic gain-of-function mutations that compensate for reduced DNMT1 protein levels. In light of these considerations, we hypothesized that DAC’s mechanistic requirements could be exploited as an activity-based selection to enrich for distal gain-of-function mutations that alter DNMT1 allostery and enhance its activity.

**Figure 1.**
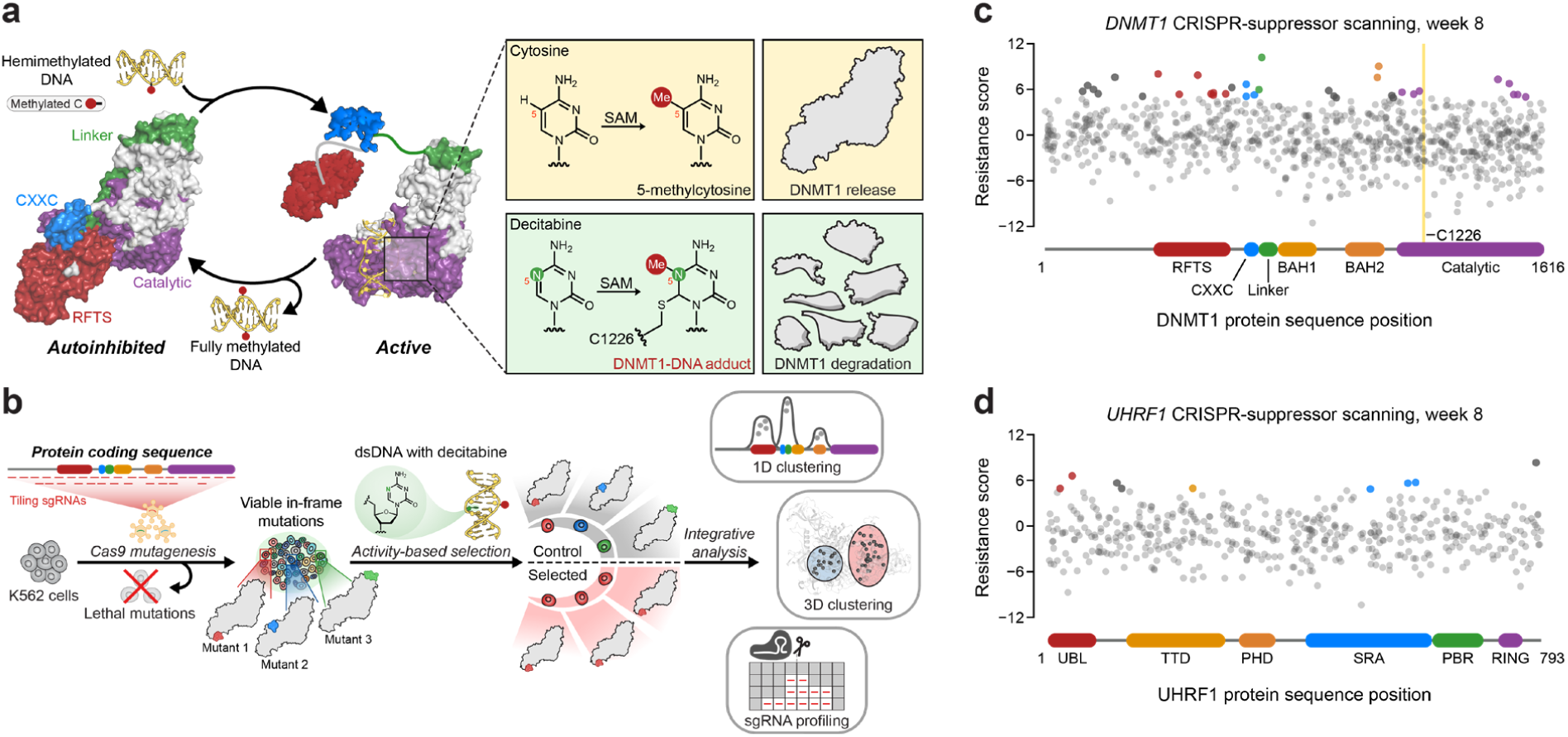
Activity-based CRISPR scanning of *DNMT1* and *UHRF1*. **a**, Schematic showing surface representations of the autoinhibited (PDB: 4WXX) and active (PDB: 4DA4) conformations of DNMT1 and DNMT1-mediated methylation of cytosine and decitabine. Methylation of decitabine leads to the formation of a covalent DNMT1–DNA adduct and subsequent proteasomal degradation. SAM, *S*-adenosyl-_L_-methionine. **b**, Schematic of the activity-based CRISPR-suppressor scanning workflow. K562 cells were transduced with an sgRNA library targeting *DNMT1* and *UHRF1* and treated with vehicle or decitabine (DAC) for 8 weeks. DAC treatment was performed at 100 nM for 5 weeks, followed by 1 μM for 3 weeks. Genomic DNA was isolated from DAC-and vehicle-treated cells and sequenced to determine sgRNA abundance. **c,d**, Scatter plots showing resistance scores (*y*-axis) for sgRNAs targeting *DNMT1* (**c**, *n* = 830) and *UHRF1* (**d**, *n* = 475) in K562 cells under DAC treatment at week 8. Resistance scores were calculated as the log_2_(fold-change) of sgRNA frequencies in DAC versus vehicle treatment, followed by normalization to the mean of the negative control sgRNAs (*n* = 77). The sgRNAs are arrayed by amino acid position in the *DNMT1* and *UHRF1* coding sequences (*x*-axis) corresponding to the positions of their predicted cut sites. Protein domains are demarcated by the colored bars along the *x*-axis. The yellow panel in **c** demarcates the position of the catalytic cysteine (C1226) in the DNMT1 active site. Data points represent the mean resistance score across three replicate treatments. Enriched sgRNAs with resistance scores greater than 2 standard deviations (s.d.) above the mean of the negative control sgRNAs are colored by their corresponding domain.

We performed CRISPR scanning in K562 cells using a pooled sgRNA library targeting the *DNMT1* and *UHRF1* coding sequences (**Fig. 1b**). After lentiviral transduction of Cas9 and the sgRNA library, the cellular pool was split and treated with vehicle (DMSO) or DAC for 8 weeks. We collected genomic DNA from the surviving cells and performed targeted amplicon sequencing of the sgRNA cassette to quantify sgRNA frequencies. We then normalized sgRNA frequencies to their initial frequencies in the library and calculated ‘resistance’ scores by comparing relative sgRNA abundance in DAC versus vehicle treatment (**Fig. 1c,d, Supplementary Table 1**).

We observed the enrichment of numerous sgRNAs after prolonged DAC treatment, consistent with the emergence and expansion of drug-resistant populations. As expected, the majority of these enriched sgRNAs targeted *DNMT1* versus *UHRF1* (**Fig. 1c,d**). Many top enriched *DNMT1* sgRNAs targeted N-terminal regulatory domains (e.g., RFTS, CXXC, BAH2), supporting the notion that resistance mutations may arise in regions distal from the active site (**Fig. 1c**). Indeed, the top enriched sgRNA in the screen, sgD702, targeted the CXXC–BAH1 linker region.

Moreover, we also detected enriched sgRNAs targeting the UBL, TTD, and SRA domains of UHRF1, suggesting that mutations beyond the direct drug target may also confer a selective advantage to DAC (**Fig. 1d**).

As enriched sgRNAs targeted diverse regions spanning *DNMT1*, we next considered whether their enrichment might indicate mutational hotspots. First, we investigated whether *DNMT1*-targeting sgRNAs exhibited any linear clustering, or contiguous amino acid intervals displaying greater enrichment than expected by chance. In brief, we used LOESS regression^31,32^ to estimate per-residue resistance scores from sgRNA resistance scores and compared them to a simulated distribution generated by shuffling sgRNA resistance scores (**Fig. 2a**, see **Methods**). We identified three linear clusters of enriched residues in *DNMT1*, spanning amino acids (aa) 119– 147 in the N-terminus, aa518–571 in the RFTS, and aa652–701 in the CXXC and linker regions (**Fig. 2b,c**). Notably, aa518–571 reside in the C-terminal lobe of the RFTS that interfaces with the catalytic domain and aa652–701 span most of the CXXC and part of the CXXC–BAH1 linker, both of which are implicated in DNMT1 autoinhibition (**Fig. 2c**).^17,18,20^ Finally, aa119–147 reside within the disordered N-terminus, which remains structurally unresolved and poorly characterized.

**Figure 2.**
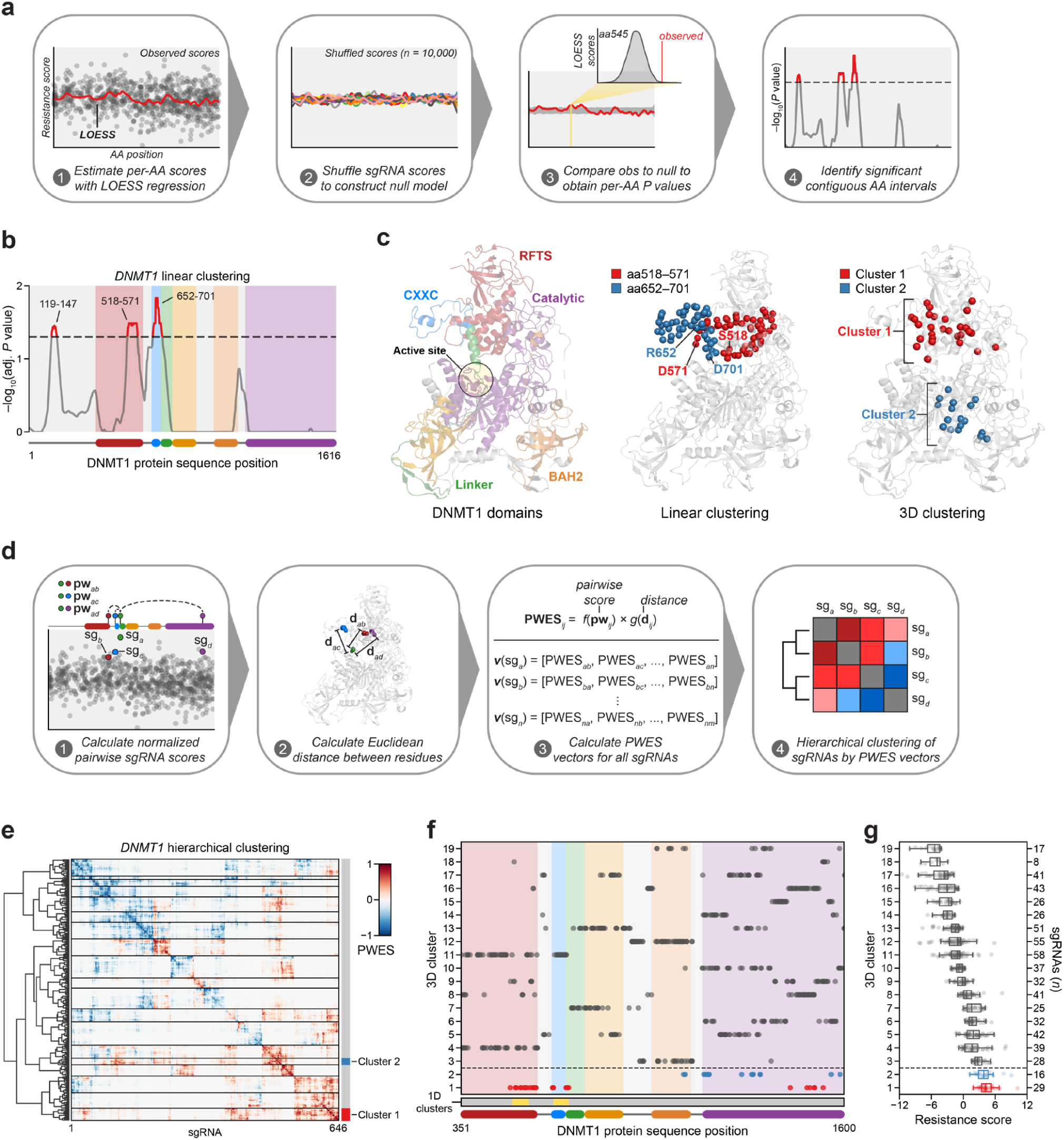
Linear and spatial clustering of CRISPR scanning data identifies putative hotspots in DNMT1 that modulate enzyme activity. **a**, Overview of the linear clustering workflow. LOESS regression and interpolation was applied to the sgRNA resistance scores to estimate per-residue resistance scores for the entire protein coding sequence. sgRNA scores were then shuffled (*n* = 10,000) and per-residue resistance scores were recalculated to simulate a null model of random enrichment. Empirical *P* values for each amino acid were determined by comparing the observed score to the simulated distribution and adjusted with the Benjamini-Hochberg procedure. Linear clusters were defined as contiguous intervals of amino acids with adjusted *P* values ≤0.05. **b**, Line plot showing −log_10_-transformed adjusted *P* values for the observed per-residue resistance scores (*y*-axis) plotted against the *DNMT1* coding sequence (*x*-axis). DNMT1 domains are demarcated by the colored background and bars along the *x*-axis. The dotted line corresponds to *P* = 0.05 and residues with *P* ≤0.05 are highlighted in red with linear clusters annotated. **c**, Structural views of autoinhibited DNMT1 (PDB: 4WXX) highlighting its domains (left panel), the linear clusters (middle panel) spanning aa518–571 (red spheres) and aa652–701 (blue spheres), and the 3D clusters 1 (red spheres) and 2 (blue spheres, right panel). The linear cluster spanning aa119–147 is not resolved in the structure. The DNMT1 active site is denoted (yellow circle) in the left panel. **d**, Overview of the 3D clustering workflow. Normalized pairwise resistance scores for sgRNAs are calculated and then scaled relative to the Euclidean distance between their targeted residues in the structural data to generate the final proximity-weighted enrichment score (PWES) for all possible pairwise sgRNA combinations. Each row or column in the resultant PWES matrix thus represents a vector of PWES values for a single sgRNA against all other resolved sgRNAs. Hierarchical clustering is applied to the PWES matrix to group sgRNAs by similarities in their overall PWES profiles. **e**, Heatmap depicting the PWES matrix of all pairwise combinations of sgRNAs (*n* = 646) targeting resolved residues in the structure of autoinhibited DNMT1 (PDB: 4WXX). sgRNAs are grouped by cluster, with black lines demarcating each cluster on the heatmap. Cluster 1 sgRNAs (*n* = 29) and cluster 2 sgRNAs (*n* = 16) are highlighted in red and blue, respectively. **f**, Scatter plot showing the targeted amino acid positions of the *DNMT1*-targeting sgRNAs used in the 3D clustering analysis. sgRNAs are grouped by 3D cluster (*y*-axis) derived from **e** and plotted against the *DNMT1* coding sequence (*x*-axis). Clusters are numbered by the mean resistance score of their component sgRNAs, with cluster 1 representing the greatest mean resistance score. Clusters 1 and 2 are highlighted in red and blue, respectively. Amino acid intervals corresponding to the linear clusters in **b** are highlighted in yellow. **g**, Box plot showing the sgRNA resistance scores (*x*-axis) for each of the 3D clusters (*y*-axis, left) derived from **e** and **f**. The number of sgRNAs per cluster are shown on the *y*-axis (right) and individual sgRNAs within each cluster are shown as dots. Clusters 1 and 2 are highlighted in red and blue, respectively. The central band, box boundaries, and whiskers represent the median, interquartile range (IQR), and 1.5 × IQR, respectively.

Functional protein regions can comprise spatially proximal residues that are distal on the linear CDS. Spatial clustering of cancer mutations at such regions is often used as evidence of protein function or positive selection.^33,34^ Therefore, we next considered whether sgRNA enrichment patterns might emerge in 3D space that are not readily observed on the linear CDS. To assess 3D sgRNA clustering, we refined and applied a structure-guided clustering methodology that we previously adapted.^14,33^ In brief, we calculated proximity-weighted enrichment scores (PWES) for all pairwise combinations of resolved sgRNAs using (1) their resistance scores and (2) the Euclidean distance between their targeted residues. Then, we performed hierarchical clustering on the resultant PWES matrix to define clusters of spatially proximal sgRNAs with similar PWES profiles (**Fig. 2d**, see **Methods**).

For 3D sgRNA-clustering, we used the structure of free, autoinhibited human DNMT1_351–1600_ (PDB: 4WXX).^20^ This structure resolves the greatest number of residues and is the only human DNMT1 structure that includes the RFTS domain, which is involved in DNMT1 autoinhibition and implicated in our 1D clustering analysis (**Fig. 2b**). We calculated PWES profiles for the 646 resolved *DNMT1*-targeting sgRNAs and performed hierarchical clustering, identifying 19 clusters of sgRNAs with varying mean resistance scores (**Fig. 2e–g, Supplementary Fig. 2a,b**). The top enriched cluster, cluster 1, comprised many sgRNAs targeting the same intervals in the RFTS, CXXC, and linker regions previously found in our 1D clustering analysis (**Fig. 2b,f**). Strikingly, these enriched cluster 1 residues span a multi-domain contact interface that is critical for mediating DNMT1 autoinhibition, suggesting that the prior linear clusters correspond to a singular 3D hotspot (**Fig. 2c**).^17,18,20^ By contrast, the second-most enriched cluster, cluster 2, mainly comprised sgRNAs targeting a region of the catalytic domain, with two additional sgRNAs targeting the BAH2 domain (**Fig. 2c**). Taken together, our findings suggest that enriched sgRNAs may target regions of DNMT1 that regulate its activity.

### Cluster 1 mutations in the RFTS, CXXC, and linker regions enhance DNMT1 activity

We next investigated cluster 1 in further detail due to its consistent enrichment across our analyses. In particular, we considered (1) the overall composition and frequency of mutations found in cluster 1 sgRNAs and (2) their functional consequences at the protein level. To assess the underlying mutational outcomes, we individually transduced K562 cells with 8 enriched cluster 1 sgRNAs targeting distinct DNMT1 regions distally positioned on the CDS. (**Fig. 3a**). Transduced cells were treated with DAC or vehicle for 8 weeks and genotyped. Raw sequencing data were processed and aligned with CRISPResso2^35^ to identify DNA variants and quantify their frequencies. We then characterized variants and their impact at the protein level with a custom pipeline (see **Methods, Supplementary Fig. 3a**). Briefly, each DNA variant was classified by the size, location, and sequence context of its mutation, and those classified as in-frame were realigned, trimmed, and translated. For downstream analyses, variants were classified into three major categories: wild-type (WT), in-frame (IF), and loss-of-function (LOF). Coding variants retaining the reference protein sequence (e.g., unedited, silent mutations) were considered wild-type. Coding mutations that maintain the reading frame, excluding nonsense and wild-type alleles, were classified as in-frame. Variants predicted to result in non-functional protein product were classified as loss-of-function.

**Figure 3.**
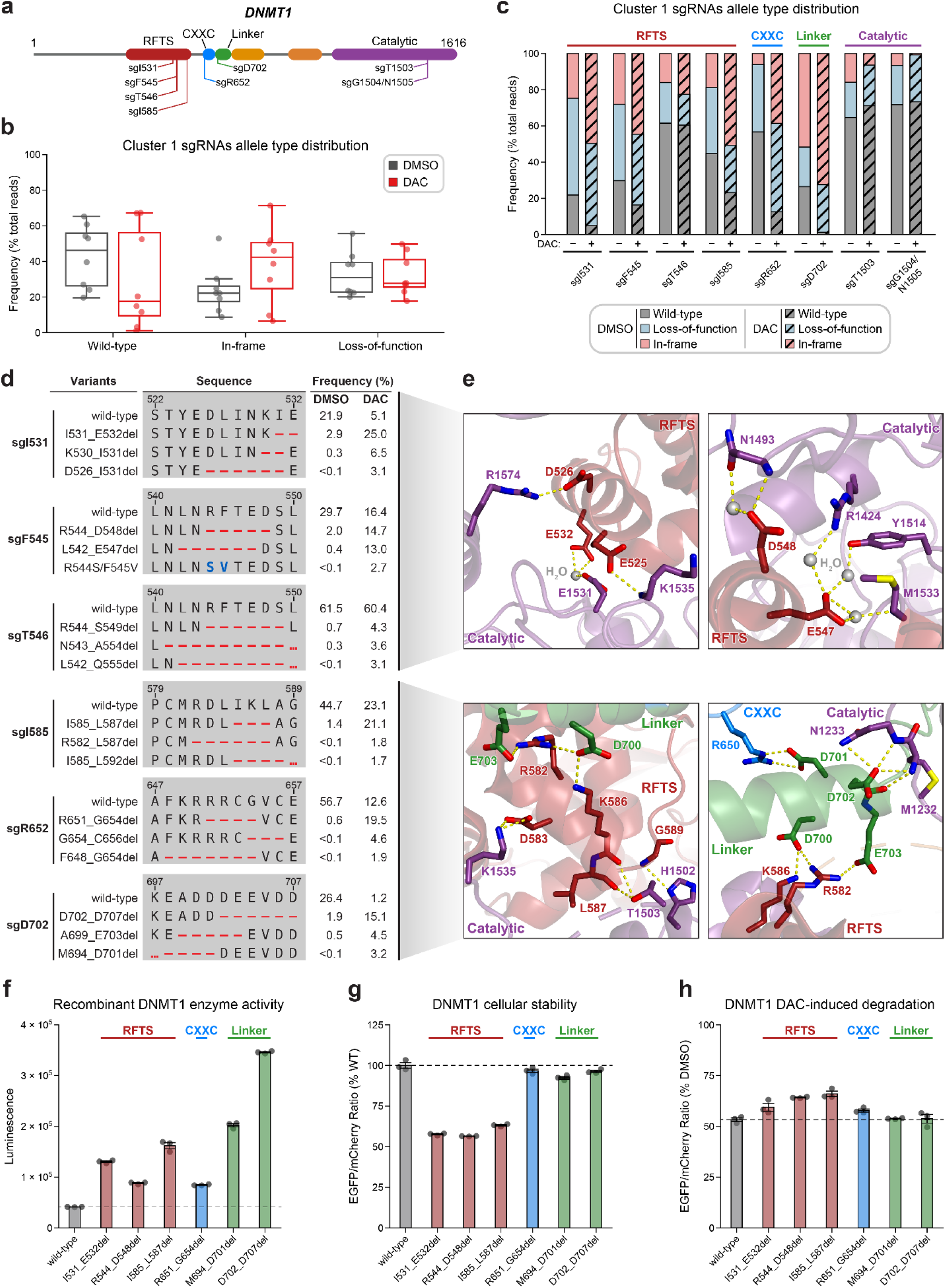
Cluster 1 sgRNAs generate hypermorphic *DNMT1* mutations in the RFTS, CXXC, and linker regions that abrogate autoinhibition. **a**, Schematic showing the amino acid positions on the *DNMT1* CDS targeted by selected cluster 1 sgRNAs (*n* = 8). **b**, Box plots showing the observed frequencies (percentage of total reads, *y*-axis) of wild-type, in-frame, and loss-of-function alleles (*x*-axis) after 8 weeks of treatment with vehicle (DMSO, gray) or 100 nM decitabine (DAC, red) for the cluster 1 sgRNAs shown in **a**. Individual sgRNAs are shown as dots. The central band, box boundaries, and whiskers represent the median, interquartile range (IQR), and 1.5 × IQR, respectively. **c**, Stacked bar plot showing the observed frequencies (percentage of total reads, *y*-axis) of wild-type, in-frame, and loss-of-function alleles for selected cluster 1 sgRNAs (*x*-axis) after 8 weeks of treatment with vehicle (DMSO) or 100 nM decitabine (DAC). **d**, Table showing the amino acid sequence alignment and observed frequencies (percentage of total reads) of the wild-type and top enriched in-frame variants across cluster 1 sgRNAs after 8 weeks of treatment with vehicle (DMSO) or 100 nM decitabine (DAC). In-frame variants were considered enriched if the observed frequency in DAC treatment was ≥1% and the log_2_(fold-change) of the observed frequency in DAC versus vehicle treatment was ≥2. Enriched in-frame variants meeting these criteria were sorted by their observed frequency in DAC treatment and the top three most abundant are shown. sgT1503 and sgG1504/N1505 were excluded due to the lack of enriched in-frame variants. Amino acid deletions are represented as red dashes and substitutions are highlighted in blue. Red ellipses are used to denote amino acid deletions that exceed the length of the shown sequence alignment. **e**, Structural views of various regions of the DNMT1 autoinhibitory interface perturbed by enriched in-frame variants (from **d**) generated by cluster 1 sgRNAs. Key residues in the RFTS (red), CXXC (blue), linker (green), and catalytic (purple) domains that mediate polar contacts are shown as sticks. Hydrogen bonds are represented by dotted yellow lines and water molecules are depicted as gray spheres. Top left, RFTS–catalytic domain interactions disrupted by sgI531. Top right, RFTS–catalytic domain interactions disrupted by sgF545 and sgT546. Bottom panels, RFTS–linker–catalytic domain interactions disrupted by sgI585, sgR652, and sgD702. **f**, Bar plot showing recombinant DNMT1 enzyme activity (luminescence, *y*-axis) for wild-type DNMT1 and selected cluster 1 variants (from **d**, *x*-axis) in the MTase-Glo assay. Wild-type is depicted in gray and variants are colored according to the domain in which the mutation is located. The dotted line represents the mean luminescence of the wild-type DNMT1 construct. Bars represent the mean ± s.d. across three replicates. One of two independent experiments is shown. **g**, Bar plot showing the cellular stability of wild-type DNMT1 and selected cluster 1 variants (*x*-axis) in K562 cells as measured by the EGFP-mCherry fluorescence reporter system. Stability was calculated as the EGFP/mCherry ratio of the construct normalized to the EGFP/mCherry ratio of wild-type DNMT1 after 3 d of vehicle treatment (percentage of wild-type EGFP/mCherry ratio, *y*-axis). The dotted line represents the mean stability of the wild-type DNMT1 construct. Bars represent the mean ± s.d. across three replicates. One of two independent experiments is shown. **h**, Bar plot showing the degradation of wild-type DNMT1 and selected cluster 1 variants (*x*-axis) in K562 cells as measured by the EGFP–IRES–mCherry fluorescence reporter system. Degradation was calculated for each construct as the EGFP/mCherry ratio of the construct in DAC treatment normalized to vehicle treatment (percentage of DMSO EGFP/mCherry ratio, *y*-axis). Cells were treated with vehicle or DAC (100 nM) for 3 d. The dotted line represents the mean degradation of the wild-type DNMT1 construct. Bars represent the mean ± s.d. across three replicates. One of two independent experiments is shown.

We observed a dramatic enrichment of in-frame mutations and concomitant depletion of the wild-type allele under DAC treatment for most cluster 1 sgRNAs (**Fig. 3b,c**), supporting the notion that in-frame mutations in cluster 1 confer a fitness advantage to DAC. We next considered whether cluster 1 in-frame variants disrupt critical interactions that mediate DNMT1 autoinhibition. To explore this possibility, we identified the top enriched in-frame variants across cluster 1 sgRNAs, excluding sgT1503 and sgG1504/N1505 due to their lack of enriched in-frame variants (**Fig. 3d, Supplementary Table 2**). Overall, enriched in-frame variants were primarily deletions ranging from −2 to −8 aa (**Supplementary Fig. 3b,c**). We first examined sgD702 as it exhibited an abundance of in-frame mutations (>70% in DAC and >50% in vehicle) and significant depletion of the wild-type allele (1.2% in DAC versus 26.4% in vehicle). Enriched in-frame mutations in sgD702 likely disrupt an α-helix in the linker, which includes a stretch of acidic residues (D700– E703) that contact the RFTS and catalytic domains to promote autoinhibition^17,18,20^ (**Fig. 3e**, bottom panels). In fact, one of the top enriched in-frame variants, M694_D701del, was identical to a previously characterized overactive DNMT1 mutant.^20^

Prompted by these observations, we sought to determine whether other enriched cluster 1 variants also structurally disrupt DNMT1 autoinhibition. Indeed, many of the top enriched variants perturbed key residues that mediate inter-domain contacts. For example, enriched in-frame mutations in the CXXC domain generated by sgR652 deleted up to 8 aa spanning F648–C656. This region contacts the linker (**Fig. 3e**, bottom right) and includes a conserved patch of basic residues (K649–R652) that when mutated has been shown to increase DNMT1 activity.^36^ Similarly, observed mutations in the RFTS domain also perturbed residues spanning three discrete intervals across this interface (D526–E532, L542–A554, M581–L592), despite their distance on the CDS. These intervals encompass α-helices that make extensive polar contacts with the linker and catalytic domains (vide infra). For sgI531, enriched in-frame variants likely compromise polar contacts with the target recognition domain (TRD), a subregion of the catalytic domain, formed by the side chains of E525, D526, and E532 (**Fig. 3e**, top left). Likewise, in-frame variants enriched in sgF545, sgT546 and sgI585 also presumably disrupt similar interactions with the TRD (E547, D548, D583) and the CXXC–BAH1 linker (R582, K586) (**Fig. 3e**). Supporting these results, prior biochemical studies have demonstrated that DNMT1 mutations at analogous or proximal positions exhibit increased methyltransferase activity compared to wild-type DNMT1.^20,36–39^ Taken together, our findings strongly suggest that cluster 1 sgRNAs may confer a selective advantage to DAC by generating gain-of-function mutations that relieve DNMT1 autoinhibition and enhance its enzymatic activity.

To confirm whether these in-frame mutations indeed enhance DNMT1 activity, we biochemically characterized a subset of enriched cluster 1 mutations in the RFTS, CXXC, and linker regions, including the previously validated overactive M694_D701del mutation (**Fig. 3d**). We purified recombinant wild-type and mutant DNMT1_351–1616_ constructs and evaluated their enzymatic activity.^39–41^ Corroborating our structural predictions, these mutants exhibited 1.7-to 5.8-fold increased activity compared to wild-type DNMT1_351–1616_ (**Fig. 3f**).

As reducing DNMT1 protein levels attenuates DNMT1–DNA adduct-induced cytotoxicity, we next considered whether cluster 1 mutations influence DNMT1 protein abundance and/or DAC-induced depletion of DNMT1.^42^ To address these questions, we evaluated cellular DNMT1 levels with a fluorescent reporter system^43,44^ in which full-length DNMT1 is fused to EGFP–IRES– mCherry. EGFP acts as a proxy for DNMT1 protein levels while mCherry serves as an internal standard, accounting for cell-to-cell variation in reporter integrations, expression, or homeostasis. We transduced K562 cells with reporter DNMT1 constructs encoding wild-type and mutant DNMT1 and assessed EGFP/mCherry fluorescence ratios after three days of treatment with DAC or vehicle. Under vehicle treatment, EGFP/mCherry ratios for the CXXC and linker mutants were comparable to wild-type DNMT1, whereas RFTS mutants displayed significantly lower EGFP/mCherry ratios, suggesting that RFTS mutations may destabilize the protein (**Fig. 3g**). These results are consistent with reports that HSAN1E and ADCA-DN patients harbor clinical *DNMT1* hotspot mutations in the RFTS domain that destabilize DNMT1.^45–47^ Upon treatment with DAC, cluster 1 mutants were degraded at similar levels as wild-type DNMT1, suggesting that these mutations are unlikely to confer a selective advantage through resistance to degradation (**Fig. 3h**). Altogether, our results collectively support a model in which cluster 1 mutations operate primarily by disrupting DNMT1 autoinhibition to enhance enzymatic activity.

### Integrative analysis reveals distinct mutational profiles between cluster 1 and 2 sgRNAs

As our approach accurately identified a validated mechanism of DNMT1 autoinhibition, we next investigated how cluster 2 sgRNAs operate. We evaluated mutational outcomes generated by 10 of the top enriched cluster 2 sgRNAs targeting the BAH2 and catalytic domains (**Fig. 4a, Supplementary Table 2**). Like before, K562 cells were individually transduced, treated with DAC or vehicle for 8 weeks, and genotyped. Notably, observed mutational outcomes were highly dissimilar between cluster 1 and cluster 2 sgRNAs. Overall, cluster 2 sgRNAs exhibited preferential enrichment of the wild-type allele and loss-of-function variants with concomitant depletion of in-frame variants, in stark contrast to the substantial enrichment of in-frame mutations seen in cluster 1 sgRNAs (**Fig. 4b**). Due to the prominent differences in their mutational profiles, we considered whether mutations found in clusters 1 and 2 operate through distinct mechanisms. Whereas cluster 1 sgRNAs induce gain-of-function mutations that enhance DNMT1’s enzymatic activity, we hypothesized that the preponderance of loss-of-function mutations found in cluster 2 sgRNAs may reduce *DNMT1* gene dosage and attenuate DAC-DNMT adduct-related cytotoxicity. Interestingly, the enrichment patterns of cluster 1 sgRNAs targeting the catalytic domain (sgT1503, sgG1504/N1505) resembled those of cluster 2 sgRNAs (**Fig. 3d**). These observations suggest that the finer genotype-level resolution of individual sgRNAs might reveal associations that are not readily apparent from screen-level data and analyses.

**Figure 4.**
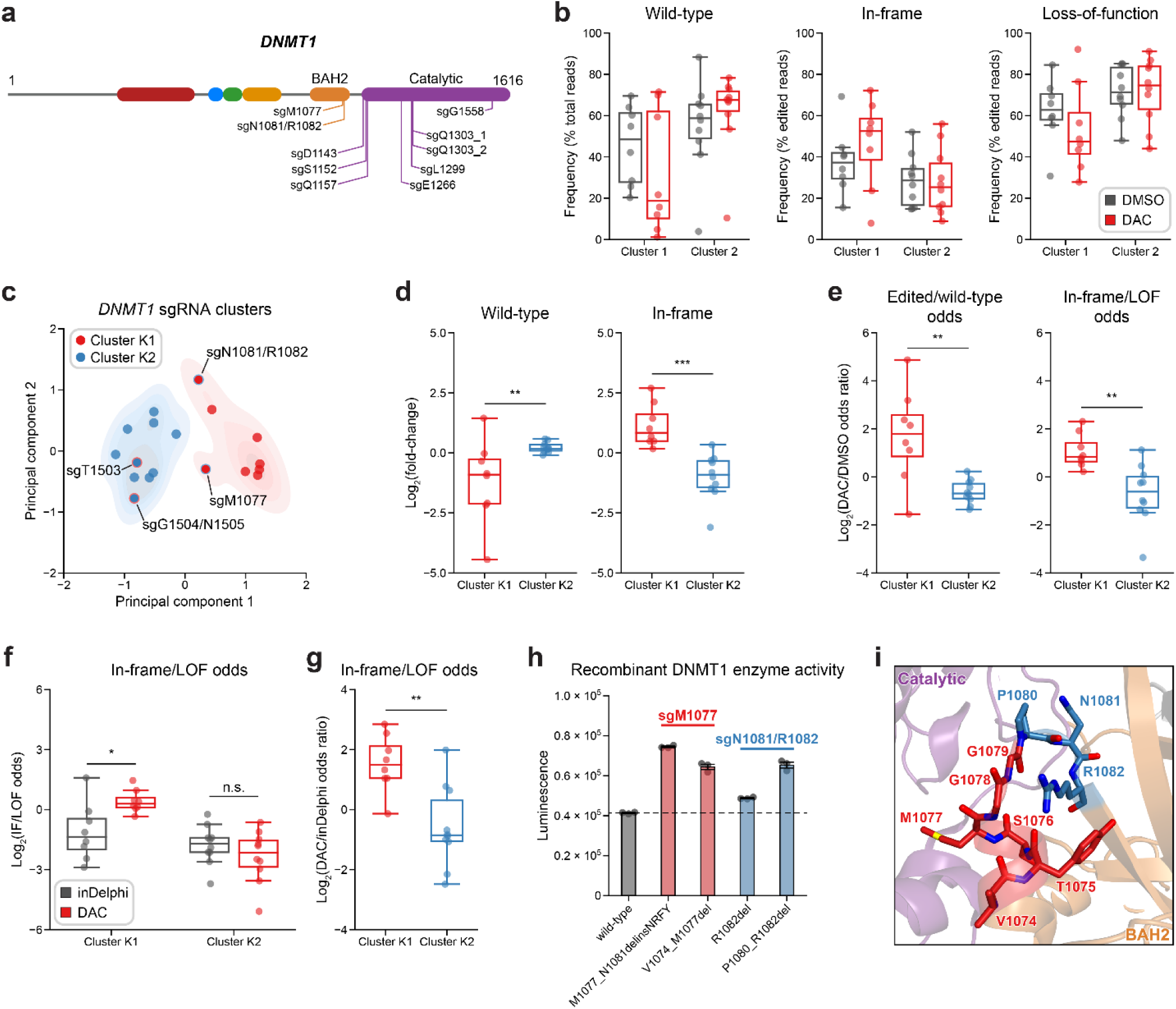
Integrative analysis reveals distinct mutational profiles between cluster 1 and 2 sgRNAs. **a**, Schematic showing the amino acid positions on the *DNMT1* coding sequence targeted by the selected cluster 2 sgRNAs (*n* = 10). **b**, Box plots showing the observed frequencies (*y*-axis) of wild-type, in-frame, and loss-of-function alleles (*x*-axis) after 8 weeks of treatment with vehicle (DMSO, gray) or 100 nM decitabine (DAC, red) for cluster 1 (*n* = 8, from **Fig. 3a**) and cluster 2 sgRNAs (*n* = 10, from **a**). Wild-type allele frequencies are plotted as the percentage of total reads. In-frame and loss-of-function allele frequencies are plotted as the percentage of edited (i.e., non-wild-type) reads. **c**, Scatter plot showing *DNMT1*-targeting sgRNAs from clusters 1 and 2 (*n* = 18) projected onto principal component space after principal component analysis (PCA) on features of their mutational profiles. The sgRNAs were then partitioned using *k*-means clustering (*k* = 2) to identify clusters K1 (red) and K2 (blue), corresponding primarily to the original 3D clusters 1 and 2. sgRNAs reassigned from cluster 1 to K2 or cluster 2 to K1 are annotated and denoted with red and blue borders, respectively. Contours depict a bivariate kernel density estimation for clusters K1 (red) and K2 (blue). **d**, Box plots showing the log_2_(fold-change) enrichment (*y*-axis) of wild-type (left) and in-frame variants (right) in DAC versus vehicle treatment for cluster K1 (*n* = 8, red) and K2 (*n* = 10, blue) sgRNAs. Log_2_(fold-change) was calculated based on the observed frequency (percentage of total reads) in DAC versus vehicle treatment. **e**, Box plots showing the log_2_(odds ratio) in DAC versus vehicle treatment (*y*-axis) for edited versus wild-type odds (left) and in-frame versus loss-of-function (LOF) odds (right) for cluster K1 (red) and K2 (blue) sgRNAs. Edited/wild-type odds were calculated using absolute frequencies (percentage of total reads) and in-frame/LOF odds were calculated using relative frequencies (percentage of edited reads). **f**, Box plots showing the log_2_(in-frame/LOF odds) for cluster K1 and K2 sgRNAs as predicted by inDelphi (gray) and as observed in DAC treatment (red). **g**, Box plots showing the log_2_(odds ratio) for in-frame/LOF odds in DAC treatment versus inDelphi predictions (*y*-axis) for cluster K1 (red) and K2 (blue) sgRNAs. **h**, Bar plot showing recombinant DNMT1 enzyme activity (luminescence, *y*-axis) for wild-type DNMT1 and selected BAH2 variants (*x*-axis) enriched in sgM1077 (red) and sgN1081/R1082 (blue) in the MTase-Glo assay. The dotted line represents the mean luminescence of the WT DNMT1 construct. Bars represent the mean ± s.d. across three replicates. One of two independent experiments is shown. **i**, Structural view of the DNMT1 BAH2 (yellow) region highlighting residues targeted by sgM1077 (red) and sgN1081/1082 (blue). The catalytic domain is shown in purple. Perturbed residues are shown as sticks and annotated. (PDB: 4WXX) For box plots in **b,d–g**, the individual sgRNAs are plotted as points and the central band, box boundaries, and whiskers represent the median, interquartile range (IQR), and 1.5 × IQR, respectively. *P* values (\**P* ≤0.05; \*\**P* ≤0.01; \*\*\**P* ≤0.001; n.s., not significant) were calculated with two-sided Mann-Whitney tests, except for **f**, where they were calculated with two-sided Wilcoxon signed-rank tests.

Subsequently, we explored whether we could distinguish individual sgRNAs through characteristics of their mutational profiles. In addition to sgRNA resistance scores, we included metrics such as the absolute frequencies (percentage of total reads) of wild-type and in-frame alleles and the relative frequency (percentage of edited reads) of in-frame alleles. Because mutational outcomes across sgRNAs can be highly complex, we also included additional features to represent the similarity and diversity of mutational outcomes. To assess mutational profile similarity in DAC versus vehicle treatment, we calculated the Pearson correlation of observed allele frequencies and the symmetric Kullback-Leibler (KL) divergence, which quantifies the similarity of two probability distributions with greater values indicating greater dissimilarity. To measure mutational diversity under DAC treatment, we calculated the Gini coefficient, a statistical measure of dispersion, with respect to all alleles and edited alleles. Finally, to capture the directionality of change across treatments, we calculated the log_2_(fold-change) of wild-type and in-frame allele frequencies, as well as the ‘odds ratio’ of edited/wild-type odds and in-frame/loss-of-function odds in DAC versus vehicle treatment. We defined the edited/wild-type odds as the proportion (percentage of total reads) of edited versus wild-type alleles and in-frame/loss-of-function odds as the proportion (percentage of edited reads) of in-frame versus loss-of-function alleles.

Using these features, we performed principal component analysis (PCA) and *k*-means clustering (*k* = 2) on the *DNMT1*-targeting sgRNAs to partition them by their mutational profiles and evaluate their similarity to our structure-derived clusters. Overall, the resultant *k*-means clusters, which we term clusters K1 and K2 to denote their resemblance to clusters 1 and 2, respectively, largely preserved the sgRNA compositions of their corresponding structure-derived clusters (**Fig. 4c**). Notably, the TRD-targeting sgRNAs (sgT1503, sgG1504/N1505) from cluster 1 were reassigned to cluster K2, in agreement with our previous observations. Conversely, our analysis also assigned the BAH2-targeting sgRNAs from cluster 2 (sgM1077, sgN1081/R1082) to cluster K1, suggesting that they share greater resemblance to core cluster 1 sgRNAs and enrich for in-frame gain-of-function variants.

We next evaluated clusters K1 and K2 across the PCA feature set to determine their distinguishing characteristics. Supporting our previous findings, cluster K1 sgRNAs displayed a greater and lower abundance of in-frame and wild-type alleles, respectively, under DAC treatment compared to cluster K2 sgRNAs (**Supplementary Fig. 4a**). Similarly, when comparing DAC versus vehicle treatment, cluster K1 sgRNAs exhibited preferential enrichment and depletion of in-frame and wild-type alleles, respectively, and greater edited/wild-type and in-frame/loss-of-function odds ratios relative to cluster K2 sgRNAs (**Fig. 4d,e**). Finally, cluster K1 mutational profiles tended to have lower Gini coefficients (all alleles) and Pearson correlations compared to cluster K2 sgRNAs, consistent with the idea that DAC-mediated positive selection of in-frame variants drives greater allelic diversity and mutational divergence relative to no selection in cluster K1 sgRNAs (**Supplementary Fig. 4b**). Taken together, our findings support the notion that cluster 1/K1 sgRNAs generate in-frame gain-of-function variants that confer a fitness advantage to DAC.

DNA sequence context significantly influences Cas9 repair outcomes.^48,49^ We therefore considered whether the observed prevalence of in-frame mutations in cluster K1 resulted from sequence biases in repair outcomes. We posited that if cluster K1 in-frame variants are under positive selection, they should be more prevalent in DAC treatment than predicted as repair outcomes. Conversely, if cluster K2 operates through a loss-of-function mechanism, then observed allele frequencies may be primarily dictated by their likelihoods as repair outcomes, as loss-of-function variants are functionally equivalent. To test this hypothesis, we used inDelphi,^48^ to predict editing outcomes from sequence context and calculated in-frame/loss-of-function odds. As expected, in-frame/loss-of-function odds were significantly greater in DAC treatment than predicted for cluster K1 sgRNAs but relatively comparable in cluster K2 sgRNAs (**Fig. 4f**). To account for baseline variations in in-frame/loss-of-function odds between sgRNAs, we calculated the odds ratio in DAC versus inDelphi and again observed significantly greater in-frame/loss-of-function odds ratios in cluster K1 versus K2 sgRNAs (**Fig. 4g**). Collectively, our results indicate that the greater prevalence of in-frame variants in DAC for cluster K1 sgRNAs stems from positive selection rather than intrinsic sequence context bias. By contrast, the comparable in-frame/loss-of-function odds between DAC and inDelphi for cluster K2 sgRNAs are consistent with a loss-of-function mechanism where observed variant frequencies are primarily linked to their probabilities as repair outcomes.

Because our analysis reassigned the cluster 2 BAH2-targeting sgRNAs to cluster K1, we considered whether these sgRNAs also select for in-frame gain-of-function variants under DAC treatment. Consequently, we functionally validated the top two enriched in-frame variants generated by sgM1077 (M1077_N1081delinsNRFY, V1074_M1077del) and sgN1081/R1082 (R1082del, P1080_R1082del) (**Supplementary Fig. 4c**). Although more modest than cluster 1 mutants, we observed 1.4-to 1.5-fold greater methyltransferase activities across the BAH2 mutants relative to wild-type, apart from R1082del, whose activity was comparable to wild-type (**Fig. 4h**). The BAH2 mutants also displayed similar levels of protein stability and DAC-induced degradation as wild-type DNMT1 in our cellular protein reporter system (**Supplementary Fig. 4d,e**). These variants target a region of the BAH2 domain near the BAH2–TRD loop, which is thought to restrain the TRD from contacting the DNA substrate in the autoinhibited conformation.^17^ Accordingly, we speculate that these BAH2 mutations may perturb mechanisms regulating substrate binding and subsequent catalysis. Altogether, our results demonstrate how genotype-level analysis can resolve mutational heterogeneity across sgRNAs within screen-derived clusters, enabling the discovery of new biological functions.

### Mutational profiles of individual sgRNAs nominate putative functional regions in UHRF1

UHRF1 is a multifunctional protein that directs DNMT1 to hemimethylated sites during DNA replication. UHRF1 is indispensable for DNA methylation maintenance, as UHRF1 ablation also causes global DNA hypomethylation.^25,26^ UHRF1-mediated recruitment of DNMT1 to chromatin requires the coordinated function of its various domains.^50,51^ Furthermore, direct and indirect interactions between UHRF1 and DNMT1 not only recruit DNMT1 to chromatin, but also stimulate its activity.^38,50–56^ Consequently, we sought to determine whether our approach could nominate gain-of-function mutations beyond the direct drug target (i.e., DNMT1). As our results demonstrate that DAC treatment enriches for hypermorphic *DNMT1* mutations, we speculated that it may also enrich for mutations in *UHRF1* that influence DNMT1 function.

Although *UHRF1*-targeting sgRNAs were enriched in our screen (**Fig. 1d**), the lack of UHRF1 structural data precluded structure-guided clustering analysis. However, our genotype-level analysis of individual sgRNAs suggests that distinct mutational characteristics can indicate functional consequences at the protein level. We therefore reasoned that this approach might identify *UHRF1*-targeting sgRNAs harboring putative gain-of-function mutations based their mutational profile signatures. Accordingly, we individually profiled 22 enriched *UHRF1*-targeting sgRNAs in K562 cells under DAC or vehicle treatment for 8 weeks like previously (**Fig. 5a, Supplementary Table 3**). We then performed PCA and *k*-means clustering analysis on the combined dataset of *UHRF1*- and *DNMT1*-targeting sgRNAs with same set of features and number of clusters as before to enable comparisons with DNMT1 clusters K1 and K2 for reference. In particular, we considered (1) how the inclusion of *UHRF1*-targeting sgRNAs might affect the partitioning of *DNMT1*-targeting sgRNAs and (2) whether *UHRF1*-targeting sgRNAs clustering with DNMT1 cluster K1 sgRNAs enrich for gain-of-function variants upon DAC treatment. Our analysis identified two comparably sized clusters (19 and 21 sgRNAs) with 11 *UHRF1*-targeting sgRNAs each (**Fig. 5b**). Reassuringly, the inclusion of *UHRF1*-targeting sgRNAs did not alter the clustering of *DNMT1*-targeting sgRNAs, indicating that these clusters may be partitioned analogously to DNMT1 clusters K1 and K2.

**Figure 5.**
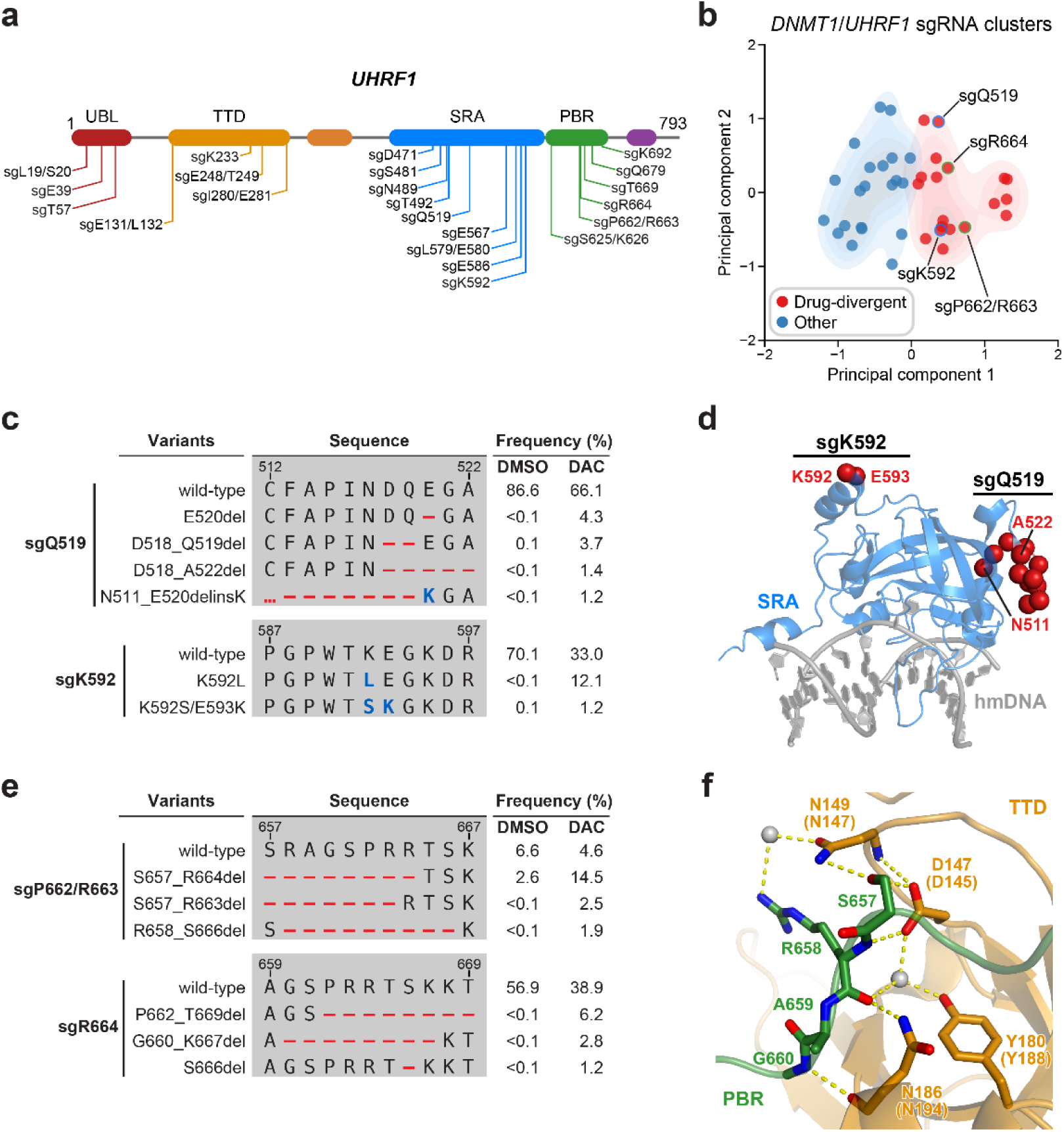
Mutational profiling analysis of individual sgRNAs nominates *UHRF1*-targeting sgRNAs with predicted gain-of-function outcomes. **a**, Schematic showing the amino acid positions on the *UHRF1* coding sequence targeted by the selected *UHRF1*-targeting sgRNAs (*n* = 22). **b**, Scatter plot showing *DNMT1*- and *UHRF1*-targeting sgRNAs (*n* = 40; 18 and 22 for *DNMT1* and *UHRF1*, respectively) projected onto principal component space after PCA on mutational profile features. The sgRNAs were then partitioned using *k*-means clustering (*k* = 2) into two categories: ‘drug-divergent’ (*n* = 19, red) or ‘other’ (*n* = 21, blue). Contours depict a bivariate kernel density estimation for drug-divergent (red) and other (blue) sgRNAs. sgRNAs highlighted in **c** and **e** targeting the SRA (blue border) and PBR (green border) regions of UHRF1 are annotated. **c**, Table showing the amino acid sequence alignment and observed frequencies (percentage of total reads) of the wild-type and top enriched in-frame variants observed in the SRA-targeting sgRNAs sgQ519 (top) and sgK592 (bottom) after 8 weeks of treatment with vehicle (DMSO) or 100 nM decitabine (DAC). In-frame variants were considered enriched if the observed frequency in DAC was ≥1% and the log_2_(fold-change) of the observed frequency in DAC versus vehicle treatment was ≥2. All enriched in-frame variants meeting these criteria are shown and ordered by their observed frequency in DAC treatment. Amino acid deletions are represented as red dashes and substitutions are highlighted in blue. Red ellipses are used to denote amino acid deletions that exceed the length of the shown sequence alignment. **d**, Structural view of the human UHRF1 SRA domain (blue) bound to hemimethylated DNA (hmDNA, gray). Residues perturbed by enriched in-frame variants (from **c**) found in sgQ519 and sgK592 are highlighted as red spheres. (PDB: 3CLZ) **e**, Table showing the amino acid sequence alignment and observed frequencies (percentage of total reads) of the wild-type and top enriched in-frame variants observed in the PBR-targeting sgRNAs sgP662/R663 (top) and sgR664 (bottom) after 8 weeks of treatment with vehicle (DMSO) or 100 nM decitabine (DAC). In-frame variants were considered enriched if the observed frequency in DAC was ≥1% and the log_2_(fold-change) of the observed frequency in DAC versus vehicle treatment was ≥2. All enriched in-frame variants meeting these criteria are shown and ordered by their observed frequency in DAC treatment. Amino acid deletions are represented as red dashes and substitutions are highlighted in blue. Red ellipses are used to denote amino acid deletions that exceed the length of the shown sequence alignment. **f**, Structural view of the zebrafish UHRF1 TTD domain (gold) complexed to a human UHRF1 PBR peptide (green) showing the region (S657–G660) targeted by sgP662/R663 and sgR664 (from **e**). Key residues forming polar contacts (yellow) are highlighted as sticks and annotated. For TTD residue annotations, the upper and lower (in parentheses) text indicate the residue identity and position in zebrafish and human UHRF1, respectively. Water molecules are shown as gray spheres. (PDB: 6B9M)

To nominate *UHRF1*-targeting sgRNAs that induce potential gain-of-function mutations, we examined the cluster containing DNMT1 cluster K1 sgRNAs, which we term ‘drug-divergent.’ As the *UHRF1*-targeting sgRNAs doubled the total number of characterized sgRNAs, we first investigated whether drug-divergent sgRNAs were analogous to DNMT1 cluster K1 sgRNAs. Indeed, the mutational profiles of drug-divergent sgRNAs shared similar characteristics to DNMT1 cluster K1 across multiple metrics (**Supplementary Fig. 5a–g**) suggesting that they may also enrich for gain-of-function variants under DAC treatment.

We next investigated drug-divergent sgRNAs targeting *UHRF1* to identify in-frame variants enriched in DAC treatment. We first examined sgQ519 and sgK592, which target the SRA domain, as they exhibited the greatest enrichment of in-frame variants and sgRNA resistance score, respectively (**Fig. 1d**). The SRA domain specifically binds hemimethylated DNA and is reported to directly interact with and stimulate DNMT1 activity.^52,57–59^ We observed substantial depletion of the wild-type allele and enrichment of multiple in-frame variants in DAC versus vehicle treatment for both sgRNAs (**Fig. 5c**). By contrast, the top 9 most prevalent mutations in sgQ519 under vehicle treatment were loss-of-function (**Supplementary Table 3**). Strikingly, the top two enriched in-frame variants for sgK592 in DAC were 3-nt and 4-nt substitutions, corresponding to K592L and K592S/E593K, respectively (**Fig. 5c**). Because point mutations are uncommon Cas9 repair outcomes^48,60^ and given their low abundance in vehicle relative to DAC treatment (K592L, 12.1% in DAC versus <0.1% in vehicle; K592S/E593K, 1.2% in DAC versus 0.1% in vehicle), we speculate that these are rare editing outcomes that are highly selected for in DAC treatment. Our previous studies have also demonstrated that such strong selection pressures can indicate stringent mutational constraints imposed by structural or functional requirements of the local sequence,^14,15^ suggesting that this region of the SRA may serve an important functional role.

Notably, enriched variants in sgQ519 and sgK592 perturb residues distal from the DNA-binding pocket of the SRA domain and the core structural elements forming the twisted β-barrel motif (**Fig. 5d**). Previous studies investigating intramolecular interactions within UHRF1 with crosslinking mass spectrometry identified extensive contacts between residues proximal to sgQ519 and sgK592 (e.g., K524, K595) and those in other UHRF1 domains.^61,62^ Accordingly, we speculate that these enriched variants are unlikely to disrupt DNA binding but may rather affect intra- or intermolecular interactions involving the SRA.

Beyond sgQ519 and sgK592, we observed several drug-divergent sgRNAs targeting the PBR region (sgP662/R663, sgR664, **Fig. 5e**), which mediates UHRF1 autoinhibition through an intramolecular interaction with the TTD domain.^63–65^ This TTD–PBR interaction maintains a ‘closed’ conformation that prevents H3K9me3 binding and recruitment to chromatin. Disrupting this interaction, such as by SRA-mediated DNA binding, drives the switch to an ‘open’ conformation that promotes H3K9me3 recognition and chromatin association. The top enriched in-frame variants induced by PBR-targeting sgRNAs perturb residues S657–G660 which are partially resolved in a structure of the zebrafish UHRF1 TTD complexed with human UHRF1 PBR peptide (**Fig. 5f**). These residues form extensive contacts with the TTD (human UHRF1 residues D145, N147, Y188, and N194), suggesting that these mutations likely disrupt the TTD–PBR interaction and promote the open conformation of UHRF1. Supporting this notion, we observed that sgK233, a drug-divergent sgRNA targeting the TTD, also generated DAC-enriched in-frame mutations disrupting key residues on the other side of the TTD–PBR interface (**Supplementary Fig. 5h,i**). Taken together, our results suggest that disrupting the TTD–PBR interaction and relieving UHRF1 autoinhibition may confer a selective advantage to DAC. More broadly, our analysis illustrates how genotype-level mutational profiling of individual sgRNAs can nominate putative gain-of-function mutations in the absence of extensive structural data.

## Discussion

Despite significant advances, the discovery of allosteric mechanisms remains challenging. Here we performed activity-based CRISPR scanning with the mechanistic inhibitor DAC to nominate multiple allosteric mechanisms regulating DNMT1 and UHRF1 function. Our study serves as an instructive framework that demonstrates how CRISPR scanning can be expanded beyond drug mechanism of action studies to identify regulatory sites in the direct drug target and even protein complex partners.

This study presents several key innovations to the CRISPR scanning methodology. First, we demonstrate how activity-based mechanistic inhibitors can enable rational screening of sophisticated phenotypes. DAC’s unique properties as an activity-based inhibitor—nearly identical to DNMT1’s native substrate—precludes the enrichment of mutations disrupting drug binding as they would likely also abrogate DNMT1 activity, which is essential for cell survival. Indeed, our results support the idea that such constraints drive the enrichment of mutations that operate through alternative resistance mechanisms, such as by enhancing DNMT1 activity. Our approach not only recapitulated mutations targeting known autoinhibitory mechanisms in DNMT1 and UHRF1, but also discovered novel hypermorphic mutations in the uncharacterized BAH2 domain of DNMT1, highlighting how activity-based CRISPR scanning can be leveraged to uncover novel biological functions. Second, our study significantly improves and expands the analysis toolkit that enables CRISPR scanning to identify putative functional hotspots. Our 1D and 3D sgRNA clustering analyses clearly implicate the DNMT1 autoinhibitory interface as a significantly enriched region of interest upon DAC treatment. Thus, our work emphasizes how an integrative approach incorporating orthogonal data, such as structural information, can nominate functional regions and provide mechanistic insights underlying their enrichment. Indeed, we expect that improvements in computational approaches and the incorporation of other data (e.g., evolutionary conservation, human genetic variation, structural predictions^66^) may further increase the power of CRISPR scanning.

Although clustering of screen-level data is effective at nominating sgRNAs and putative hotspots for further validation, our findings demonstrate how the mutational profiles of individual sgRNAs at genotype-level resolution can uncover diverse responses to drug treatment that cannot be observed with screen-level enrichment scores. By clustering individual sgRNAs using their mutational profiles, we show that DNMT1 clusters K1 and K2 largely recapitulate the 3D structure-derived clusters with notable exceptions, such as the BAH2-targeting sgRNAs in cluster 2. Through an in-depth comparative analysis, we demonstrate that cluster K1 and K2 sgRNAs exhibit unique mutational signatures upon DAC treatment and identify defining characteristics of cluster K1 sgRNAs, such as the enrichment of in-frame mutations, that may be predictive of functional consequences. To validate our approach, we biochemically characterized a subset of these enriched BAH2 variants and show that these mutations are gain-of-function and enhance DNMT1 activity. Collectively, our analysis demonstrates that the greater resolution of genotype-level data can reveal significant mutational heterogeneity across enriched sgRNAs, and that their mutational signatures can be exploited as a heuristic to nominate those with unique functional outcomes.

Finally, we showcase how this heuristic approach can be generalized to nominate sgRNAs that generate functional protein variants, especially in the absence of extensive structural information. We applied our mutational analysis methodology to nominate potential sites within UHRF1, a poorly characterized partner of DNMT1, that modulate DNMT1 function. Using a combined dataset of *UHRF1-* and *DNMT1-*targeting sgRNAs, we defined a set of ‘drug-divergent’ sgRNAs with similar characteristics as DNMT1 cluster K1 to identify putative gain-of-function variants in UHRF1. Strikingly, drug-divergent sgRNAs targeting *UHRF1* enriched for in-frame variants that perturb key residues on both sides of the TTD–PBR interface that mediates UHRF1 autoinhibition, in addition to uncharacterized regions of the SRA domain. While further study is necessary to elucidate how these UHRF1 mutations may affect DNMT1 activity, our findings outline how evaluating enriched sgRNAs at the genotype level can nominate variants for further functional validation despite a lack of structural data.

Altogether, here we demonstrate how activity-based CRISPR scanning can be leveraged to nominate allosteric regulatory mechanisms, using DNMT1 and UHRF1 as an instructive paradigm. Through an array of genetic, biochemical, and computational approaches, we illustrate how integrative analyses can offer mechanistic insights at increasing levels of resolution. In summary, our study establishes a framework for applying CRISPR scanning to systematically identify allosteric mechanisms and other complex phenotypes across various protein targets.

## Data Availability

Data are provided in the main text and figures, supplementary figures (**Supplementary Figures 1–5**) and tables (**Supplementary Tables 1–4**).

## Code Availability

Custom code used in this study will be made available on GitHub prior to publication.

## Acknowledgements

We thank members of the Liau Lab for helpful discussions and comments on the manuscript, in particular A. Siegenfeld and A. Waterbury. We thank K. Zhao and P. Randolph for assistance with next-generation sequencing. We thank E. Dolen and R. Switzer for advice regarding protein expression and purification. K.C.N., N.Z.L., and E.M.G. were supported by NSF Graduate Research Fellowships (grant no. DGE1745303). E.M.G. was also supported by the Landry Cancer Biology Fellowship. C.L. was supported by the Herchel Smith Graduate Fellowship. This research was supported by award no. 1DP2GM137494 from the National Institute of General Medical Sciences, the Damon Runyon-Rachleff Innovation Award, and startup funds from Harvard University.

## Author Contributions

K.C.N. and B.B.L. conceived the study, designed experiments, and wrote the manuscript. K.C.N. and S.M.H performed experiments with assistance from P.M.G., D.A.T., N.Z.L., E.G., and C.L. K.C.N analyzed data with assistance from S.M.H., D.A.T., and C.L. All authors edited and approved the manuscript. B.B.L. supervised and held overall responsibility for the study.

## Methods

### Chemical reagents

Compounds were stored at −80 °C in 100% DMSO (Sigma-Aldrich). The vehicle condition represents 0.1% DMSO treatment. Decitabine (DAC) was purchased from Selleck Chemicals (≥99% purity by HPLC).

### Cell culture

K562 cells were obtained from ATCC. HEK293T cells were a gift from B.E. Bernstein (Massachusetts General Hospital). All cell lines were cultured in a humidified 5% CO_2_ incubator at 37 °C and routinely tested for mycoplasma (Sigma-Aldrich). All media were supplemented with 100 U ml^−1^ penicillin and 100 µg ml^−1^ streptomycin (Gibco) and fetal bovine serum (FBS, Peak Serum). K562 cells were cultured in RPMI-1640 (Gibco) supplemented with 10% FBS. HEK293T cells were cultured in DMEM (Gibco) supplemented with 10% FBS.

### Lentivirus production and transduction

To produce lentivirus, transfer plasmids were co-transfected with *GAG/POL* and *VSVG* plasmids into HEK293T cells using FuGENE HD (Promega) and the medium was exchanged 6–8 h after transfection. Viral supernatant was collected 48–60 h after transfection, filtered (0.45 µm), and stored at −80 °C until use. K562 cells were transduced by spinfection at 1,800*g* for 90 min at 37 °C with 8 µg ml^-1^ polybrene (Santa Cruz Biotechnology) and selected with 2 µg ml^-1^ puromycin (Gibco) starting 48 h post-transduction.

### Plasmid construction

For individual sgRNA validation experiments, single-stranded oligonucleotides encoding the sgRNA (Sigma-Aldrich) were annealed and cloned into lentiCRISPRv2 (a gift from F. Zhang, Addgene #52961) using Golden Gate cloning with FastDigest Esp3I (Thermo Fisher Scientific) and T4 Ligase (New England Biolabs). For protein cellular stability and degradation experiments, full-length human DNMT1 was cloned into the Artichoke reporter plasmid (a gift from B. Ebert, Addgene #73320) using Gibson Assembly with NEBuilder HiFi (New England Biolabs). For bacterial expression constructs, truncated human DNMT1 (residues 351–1616) was cloned into pET15b containing an N-terminal His_6_-tag and TEV protease cleavage site using Gibson Assembly. The wild-type human DNMT1 CDS was subcloned from pcDNA3.1–HA–DNMT1, a gift from D. Tenen (Harvard Medical School) and mutations were introduced by modifying the primers used to amplify the DNMT1 CDS for Gibson Assembly.

### Cellular protein stability and degradation assays

Lentivirus was produced for wild-type and mutant DNMT1–EGFP–IRES–mCherry (Artichoke) reporter constructs as described above. Wild-type K562 cells were then transduced with the appropriate lentivirus and selected with puromycin for 3 d as described above. The selected cells for each construct were then split into two pools and treated with 100 nM DAC or vehicle for 3 days in triplicate, after which EGFP and mCherry fluorescence were measured on a NovoCyte 3000RYB flow cytometer (Agilent). The geometric mean of the ratio of EGFP to mCherry fluorescence was calculated for mCherry-positive cells (see **Supplementary Fig. 3d**) in each sample using the NovoExpress software (Agilent). To assess cellular stability, the EGFP/mCherry ratios of the mutant constructs in vehicle treatment were normalized to the wild-type EGFP/mCherry ratio in vehicle treatment. To assess degradation, the EGFP/mCherry ratios in DAC treatment for each construct were normalized to their respective EGFP/mCherry ratios in vehicle treatment.

### Immunoblotting

Cells were treated with vehicle or varying concentrations of DAC for 48 h and then harvested. Cells were washed three times with cold PBS (Corning) and lysed in RIPA buffer (Boston BioProducts) supplemented with 1× Halt Protease Inhibitor Cocktail and 5 mM EDTA (Thermo Fisher Scientific) on ice for 30 min. Lysates were clarified by centrifugation and total protein concentration in clarified lysates was determined using the BCA Protein Assay Kit (Thermo Fisher Scientific) prior to preparing samples for SDS-PAGE. Immunoblotting was performed according to standard procedures using the following primary antibodies: anti-DNMT1 (clone D63A6, Cell Signaling Technology #5032, 1:1,000) and anti-LMNB1 (Abcam #ab16048, 1:2,000).

### Protein expression and purification

Wild-type and mutant human DNMT1_351–1616_ bacterial expression constructs were cloned as described above. Recombinant DNMT1 expression and purification was performed according to published protocol^39^ with some modifications. DNMT1 expression constructs were transformed and expressed recombinantly in *Escherichia coli* Rosetta2(DE3)pLysS cells (Novagen). Freshly transformed cells were grown in LB broth supplemented with ampicillin and chloramphenicol at 37 °C to an OD_600_ of 0.6, after which the cells were cooled on ice and induced with 0.4 mM isopropyl-β-D-thiogalactoside (IPTG, Research Products International) at 16 °C overnight. Cells were harvested, pelleted by centrifugation, and stored at −80 °C until use. Cells were resuspended in lysis buffer containing 25 mM Tris-HCl (pH 7.5), 500 mM NaCl, 4 mM β-mercaptoethanol (BME), 5% glycerol, 3 U ml^-1^ DNase I (New England Biolabs), and 1× cOmplete EDTA-free protease inhibitor cocktail (Roche), lysed by sonication, and clarified by centrifugation. Clarified lysate was incubated with His60 Ni Superflow resin (Takara Bio) for 1 h at 4 °C and then washed with buffer containing 20 mM Tris-HCl (pH 7.5), 500 mM NaCl, 4 mM BME, 5% glycerol, and 20 mM imidazole. Protein was eluted with buffer containing 20 mM Tris-HCl (pH 7.5), 500 mM NaCl, 4 mM BME, 5% glycerol, and 400 mM imidazole. The eluate was diluted with an equal volume of buffer containing 20 mM sodium phosphate (pH 7.5), 2 mM dithiothreitol (DTT), and 5% glycerol and then further purified on a HiTrap Heparin HP column (Cytiva) using a linear gradient of 0.25–1.5 M NaCl in buffer containing 20 mM sodium phosphate (pH 7.5), 2 mM DTT, and 5% glycerol. Fractions containing DNMT1 were pooled and concentrated with Amicon Ultra 30 kDa centrifugal filters (EMD Millipore) and purity was verified by SDS-PAGE. Purified proteins were quantified by absorbance at 280 nm and stored in 40% glycerol at −80 °C until use.

### DNMT1 enzymatic activity assays

DNMT1 enzymatic activity was measured using the MTase-Glo Methyltransferase Assay (Promega) with recombinant DNMT1_351–1616_ and a 14-bp oligonucleotide substrate (Integrated DNA Technologies) containing a single hemimethylated CpG site. Methyltransferase reactions were prepared with 800 nM recombinant DNMT1_351–1616_, 10 μM hemimethylated DNA substrate, 10 μM *S*-adenosyl-L-methionine (SAM), and 1× MTase-Glo Reagent in reaction buffer (20 mM Tris-HCl (pH 8.0), 50 mM NaCl, 3mM MgCl_2_, 1 mM DTT, 1 mM EDTA, 0.1 mg ml^-1^ BSA) and incubated for 90 min at 30 °C. Following incubation, an equal volume of MTase-Glo Detection Solution was added to each reaction and further incubated for 30 min at room temperature. Reactions were then plated in technical triplicate in 20 μl volumes into a white, opaque 384-well plate (Corning) and endpoint luminescence was measured using a SpectraMax i3x plate reader (Molecular Devices). To account for baseline luminescence, raw luminescence values were corrected by subtracting the average luminescence values of control reactions prepared without DNMT1. All activity assays were independently conducted at least twice.

### Pooled sgRNA library cloning and CRISPR scanning experiments

The pooled sgRNA tiling library was designed with CRISPOR^67^ using the following criteria: (1) the 20-nt protospacer sequence must be upstream of an NGG PAM, (2) the predicted cleavage site falls within the coding sequence of *DNMT1* (*NP_001370*.*1*) or *UHRF1* (*NP_001041666*.*1*), (3) the sgRNA must have an off-target score (MIT Specificity Score) greater than 20. All sgRNAs meeting these criteria were synthesized as an oligonucleotide pool (Twist Biosciences) and their sequences are listed in **Supplementary Table 1**. The sgRNA oligo pool was amplified, cloned into lentiCRISPRv2, and sequenced to confirm sgRNA representation as previously described.^68,69^ Lentivirus carrying the resulting pooled sgRNA tiling library was produced as described above and titered according to published procedure.^68,69^

For CRISPR scanning, K562 cells (40 × 10^6^) were transduced at a multiplicity of infection <0.3 and subsequently selected with puromycin for 4 d. Cells were then split into pools and treated with DAC (100 nM for 5 weeks and then 1 μM for 3 weeks) or vehicle in triplicate. The cells were passaged every 3–4 d at a seeding density of 0.1–0.2 × 10^6^ cells ml^-1^ into fresh media containing drug or vehicle. Genomic DNA was isolated using the QIAamp DNA Blood Mini Kit (Qiagen). To measure the sgRNA composition of the population, the sgRNA expression cassette was PCR amplified using barcoded primers, purified, and sequenced as previously described.^14,68,69^ All samples were sequenced on a MiSeq (Illumina) using 150-cycle, single-end reads. Sufficient coverage of the sgRNA library was maintained in accordance with published recommendations.^68,69^

### CRISPR scanning data analysis

All data processing and analysis were performed using Python v.3.8.3 (www.python.org). Raw sequencing data were processed as previously described.^14,15^ In brief, reads were counted by identifying the 20-nt sequence downstream of the ‘CGAAACACCG’ prefix and mapped against a reference file containing all library sgRNA sequences with no mismatch allowance. sgRNAs with zero reads in the plasmid library were excluded from the analysis. Read counts were then converted to reads per million, increased by a pseudocount of 1, log_2_-transformed, and then normalized by subtracting the log_2_-transformed sgRNA counts in the plasmid library. Library-normalized scores were averaged across replicates for each condition and sgRNA ‘resistance scores’ were calculated by first subtracting the scores in vehicle treatment from their corresponding scores in DAC treatment, and then further normalized by subtracting the mean resistance score of the negative control sgRNAs from all sgRNAs. sgRNAs were classified as ‘enriched’ if their resistance scores were greater than the mean resistance score plus two standard deviations of the negative control sgRNAs. sgRNAs were assigned to protein amino acid positions by using the genomic coordinates of their predicted cut sites in the *DNMT1* (*NP_001370*.*1*) or *UHRF1* (*NP_001041666*.*1*) coding sequences. sgRNAs were assigned to a single amino acid if the cut site fell within a codon or assigned to the two flanking amino acids if the cut site fell between codons.

### Linear clustering analysis

As proximal sgRNAs can exhibit significant variation due to sgRNA-specific factors (e.g., off-target activity, cutting efficiency), per-residue resistance scores were estimated with respect to local sgRNAs by using LOESS regression to fit the observed sgRNA resistance scores as a function of amino acid position. To estimate resistance scores for each amino acid in DNMT1, LOESS regression was performed using the ‘lowess’ function of the statsmodels package (v.0.12.1) in Python with a 100 AA sliding window (‘frac = (100 AA/*L*)’, where *L* is the total length of the protein), and ‘it = 0’. For amino acid positions that were not targeted by sgRNAs, resistance scores were interpolated by performing quadratic spline interpolation on the LOESS output scores using the ‘interp1d’ function of the SciPy^70^ package (v.1.7.1).

To assess statistical significance of the resulting clusters, we simulated a null model of random sgRNA enrichment. sgRNA cut site positions were kept fixed while sgRNA resistance scores were randomly shuffled, and per-residue resistance scores were recalculated by performing LOESS regression and interpolation on the randomized sgRNA resistance scores for each of 10,000 permutations. Empirical *P* values were calculated for each amino acid by comparing its observed resistance score to the null distribution of random resistance scores. Empirical *P* values were adjusted using the Benjamini-Hochberg procedure to control the false discovery rate (FDR) to

≤0.05. Finally, linear clusters were called by identifying all contiguous intervals of amino acids with adjusted *P* values ≤0.05.

### 3D spatial clustering analysis

To perform the 3D spatial clustering analysis, we first calculated proximity-weighted enrichment scores (PWES) between pairs of sgRNAs. We employed a modified version of previously published procedures,^14,33^ using a scoring function involving (1) the pairwise score for a given pair of sgRNAs and (2) the Euclidean distance between their targeted residues. First, for all combinations of *DNMT1*-targeting sgRNAs *i* and *j*, the pairwise score pw_*i,j*_ was calculated using the following function:

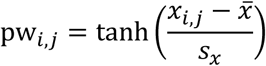

where *x*_*i,j*_ is the sum of the sgRNA resistance scores for a given pair of sgRNAs *i* and *j* (*i* ≠’ *j*), while 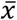 and *s*_*x*_ are the mean and standard deviation, respectively, of the summed resistance scores for all pairwise combinations of *DNMT1*-targeting sgRNAs. The hyperbolic tangent function was used to scale pairwise scores in order to minimize the disproportionate influence of highly enriched or depleted (i.e., jackpotted) sgRNAs and normalize them into the interval of [−1,1].

Next, we determined the distances between all pairwise combinations of resolved amino acids in the structure of human DNMT1_351–1600_ (PDB: 4WXX) by calculating the Euclidean distance between the centroids of the two residues using PyMOL (v.2.5.0, Schrödinger). We then isolated the subset of sgRNAs whose assigned amino acid positions were resolved in the structure. sgRNAs predicted to cut between residues were assigned to the even-numbered residue. Thus, the final PWES for all pairwise combinations of resolved sgRNAs *i* and *j* were calculated as follows:

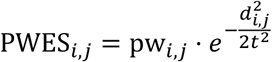

where *d*_*i,j*_ is the Euclidean distance between the targeted residues of sgRNAs *i* and *j* and *t* = 16. Hierarchical clustering was performed as previously described on the resultant pairwise PWES matrix to group sgRNAs by their PWES profiles.^14^

To assess the significance of clusters 1 and 2, their sgRNA resistance scores were kept fixed while their targeted amino acid positions were randomly shuffled (*n* = 10,000) across the amino acids targeted by resolved sgRNAs to simulate a null distribution of PWES values with randomized spatial proximity. We then took the sum of the absolute values of PWES for all intra-cluster pairwise sgRNA combinations to calculate the ‘summed PWES’ score per cluster. Empirical *P* values were calculated by comparing the observed summed PWES score for a cluster to the simulated distribution of summed PWES scores.

### Individual sgRNA validation experiments and genotyping data analysis

Individual sgRNAs selected for further validation were cloned into lentiCRISPRv2 as described above. The sequences for the sgRNAs used in these experiments are listed in **Supplementary Table 4**. Lentivirus was produced separately for each sgRNA construct as described above and K562 cells were transduced and selected with puromycin for 5 d. For each sgRNA transduction, the selected cells were then split into two pools and treated with 100 nM DAC or vehicle for 8 weeks. Genomic DNA was isolated from harvested cells using QuickExtract DNA Extraction Solution (Lucigen) and used to prepare libraries for next-generation sequencing as described previously.^68^ Briefly, the genomic region surrounding the predicted cut site of each sgRNA was first PCR amplified using genomic primers with Illumina adapters, followed by a second round of PCR to attach barcodes to the final amplicons. The final amplicons were then gel-purified using the Zymoclean Gel DNA Recovery Kit (Zymo Research), pooled, and sequenced on an Illumina MiSeq using 300-cycle, single-end reads. Primer sequences are provided in **Supplementary Table 4**.

To identify genomic variants and quantify allele frequencies, raw sequencing data were processed and aligned to *DNMT1* and *UHRF1* using CRISPResso2^35^ (v.2.0.40) with the following parameters: ‘-w 30 -q 10 –min_bp_quality_or_N 10 –exclude_bp_from_left 5 – exclude_bp_from_right 5 –plot_window_size 30’. CRISPResso2 allele frequency outputs were further processed with custom Python scripts to classify and characterize variants at the protein level for downstream analysis. Reads with no editing were classified as ‘wild-type.’ For all variants with mutations within the coding sequence, variants were classified as ‘in-frame’ if the net indel size was a multiple of three and ‘frameshift’ if not. Variants with mutations that span an intron-exon junction or otherwise disrupted canonical splice site positions (the 2 nt immediately flanking each exon) were classified as ‘splice site disrupting.’ In-frame variants were then further processed into their corresponding protein variants by performing global re-alignment to the reference CDS at the nucleotide level using a customized codon-based implementation of the Needleman-Wunsch algorithm using the ‘PairwiseAligner’ module of Biopython^71^ (v.1.7.8), followed by trimming and translation. Translated in-frame variants were further classified as ‘nonsense’ if the mutation led to a premature stop codon or merged with ‘wild-type’ if the translation protein variant matched the reference protein sequence (e.g., silent mutations, SNPs). Finally, all variants identified as frameshift, splice site disrupting, or nonsense were further classified as ‘loss-of-function.’ After processing, the final allele tables for each sgRNA were filtered to only include variants with read frequencies ≥0.1% in either DAC or vehicle treatment and frequencies re-normalized to 100%. Processed variants and their read frequencies are supplied in **Supplementary Tables 2 and 3**.

Editing outcome predictions for individual gRNAs were obtained using the inDelphi^48^ web server (https://indelphi.giffordlab.mit.edu) in single mode with K562 as the cell type. As inDelphi does not consider intron-exon boundaries or translated protein products, inDelphi-predicted genotypes were also processed similarly as above in order to classify variants and accurately determine the predicted frequencies of in-frame versus loss-of-function mutations.

### Individual sgRNA mutational profile analysis and clustering

Processed and filtered allele tables were used to calculate the various metrics for the mutational profile analysis of individual sgRNAs. Absolute variant frequencies were calculated by dividing the reads assigned to a particular variant by the total number of reads. Relative variant frequencies were calculated with respect to the total number of reads assigned to edited (i.e., in-frame or loss-of-function, wild-type excluded) variants. Log_2_(fold-change) metrics for wild-type, in-frame, and loss-of-function mutation types were calculated as the absolute frequency of the mutation type in DAC divided by the absolute frequency in vehicle, followed by log_2_-transformation.

Log-odds were calculated for the two binary outcomes of edited versus wild-type (edited/WT) and in-frame versus loss-of-function (IF/LOF) as follows:

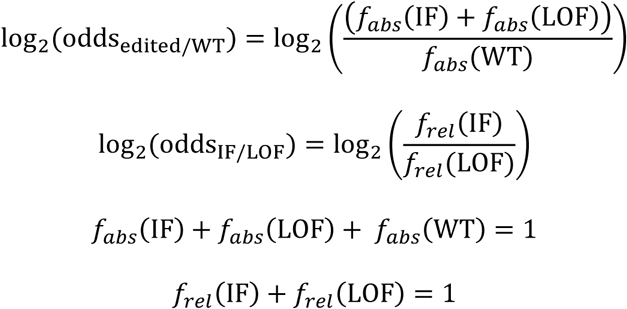

where *f*_*abs*_ and *f*_*rel*_ refer to the absolute and relative frequency, respectively, of the mutation type. Log-odds for edited versus wild-type alleles were calculated using absolute frequencies, whereas log-odds for in-frame versus loss-of-function alleles were calculated using relative frequencies. Log-odds ratios comparing DAC to vehicle or inDelphi were calculated by subtracting the log-odds in vehicle or inDelphi from the log-odds value in DAC.

Pearson correlations were calculated on the absolute variant frequencies in DAC and vehicle treatments using the ‘stats.pearsonr’ function of SciPy.^70^ Gini coefficients were calculated as follows:

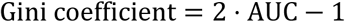

where AUC is the area under the curve of the empirical cumulative distribution function of allele frequencies. Gini coefficients were calculated with respect to all alleles using absolute frequencies as well as edited alleles using relative frequencies.

To assess the similarity of mutational profiles across treatment conditions, we used the symmetric Kullback-Leibler (KL) divergence, which is calculated for two probability distributions *P* and *Q* (i.e., allele frequency distributions in DAC and vehicle, respectively) as the sum of the standard KL divergences of *P* from *Q* and *Q* from *P* as follows:

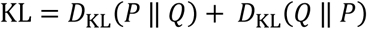

where the standard KL divergence of *P* from *Q* is calculated as:

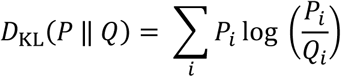

where *i* indexes the alleles found in each sgRNA, and *P*_*i*_ and *Q*_*i*_ are the frequencies of allele *i* in samples *P* and *Q* (i.e., DAC and vehicle treatments). To avoid division by zero, a pseudocount of 0.01% was added to all allele frequencies.

Subsequent data preprocessing, principal component analysis (PCA), and *k-*means clustering were performed in Python using the scikit-learn package (v.0.24.2).^72^ Feature input data were first preprocessed by independently applying a rank-based quantile transformation on each feature using the ‘QuantileTransformer’ function. PCA was performed on the transformed dataset using the ‘sklearn.decomposition.PCA’ function with ‘n_components = 10’. To cluster sgRNAs based on their mutational profile features, *k*-means clustering was performed on the resultant PCA matrix using the ‘sklearn.cluster.KMeans’ function with ‘n_clusters = 2’ and ‘n_init = 1000’. To verify the fidelity of the clusters, *k*-means clustering was performed 1,000 times and the most common outcome was used for the final *k*-means cluster assignments.

### Statistical methods and replication

Statistical parameters including the exact value and definition of *n*, the definition of center, dispersion, precision measures (mean ± s.d. or s.e.m) and statistical significance are reported in figures and figure legends. All statistical tests were performed as two-sided tests using the SciPy package (v1.7.1). All experiments were performed at least twice except for those involving next-generation sequencing, which were conducted once.

## Supplementary Figures

**Supplementary Figure 1.**
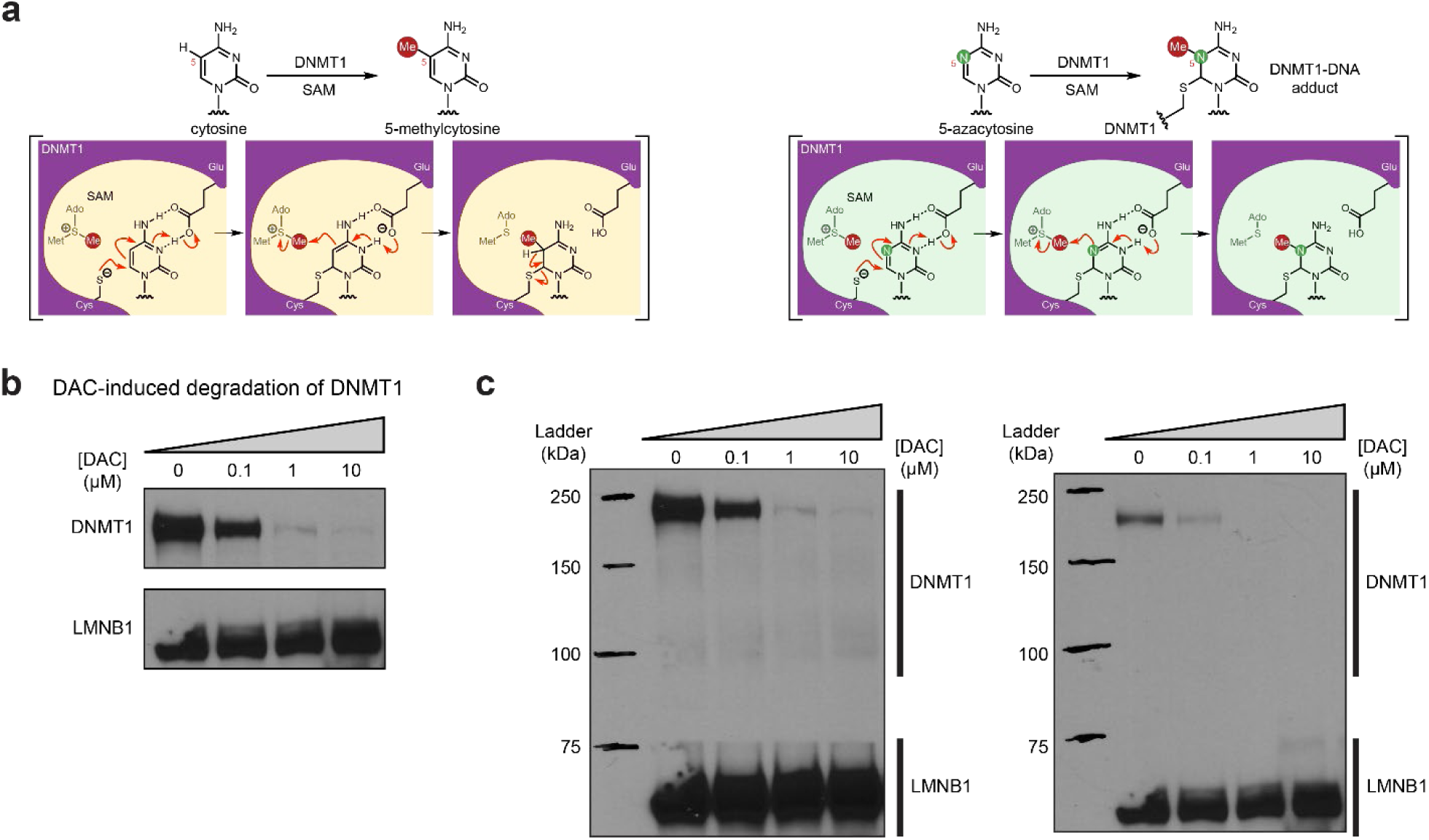
Activity-based CRISPR scanning of *DNMT1* and *UHRF1*. **a**, Schematic showing key intermediate steps in the mechanism of DNMT1-catalyzed methylation of cytosine (left) and 5-azacytosine (right), the cytosine analog found in decitabine (DAC). Nucleophilic attack on cytosine C6 by the catalytic cysteine (Cys1226 in human DNMT1) leads to the formation of a DNMT1-DNA covalent intermediate. Deprotonation of N3 by a conserved glutamic acid residue (Glu1266 in human DNMT1) facilitates the transfer of the methyl group from *S*-adenosyl-L-methionine (SAM) to the cytosine C5 position. Finally, deprotonation at the C5 position leads to β- elimination of the thiolate, releasing DNMT1 from the DNA. Conversely, the replacement of C5 with a nitrogen in the 5-azacytosine moiety of decitabine prevents this final β-elimination step and dissociation of DNMT1, thus leading to the formation of a covalent DNMT1–DNA adduct. **b**, Immunoblot showing DAC-induced degradation of DNMT1 in K562 cells after 48 hours of treatment at the indicated concentrations of DAC. One of two independent experiments is shown. The loading control (LMNB1) shown is taken from a separate exposure of the same blot. Uncropped images of the two exposures are shown in **c**. **c**, Uncropped images of the immunoblots shown in **b**. Bands for DNMT1 were taken from the exposure on the left. The loading control (LMNB1) bands were taken from the exposure on the right.

**Supplementary Figure 2.**
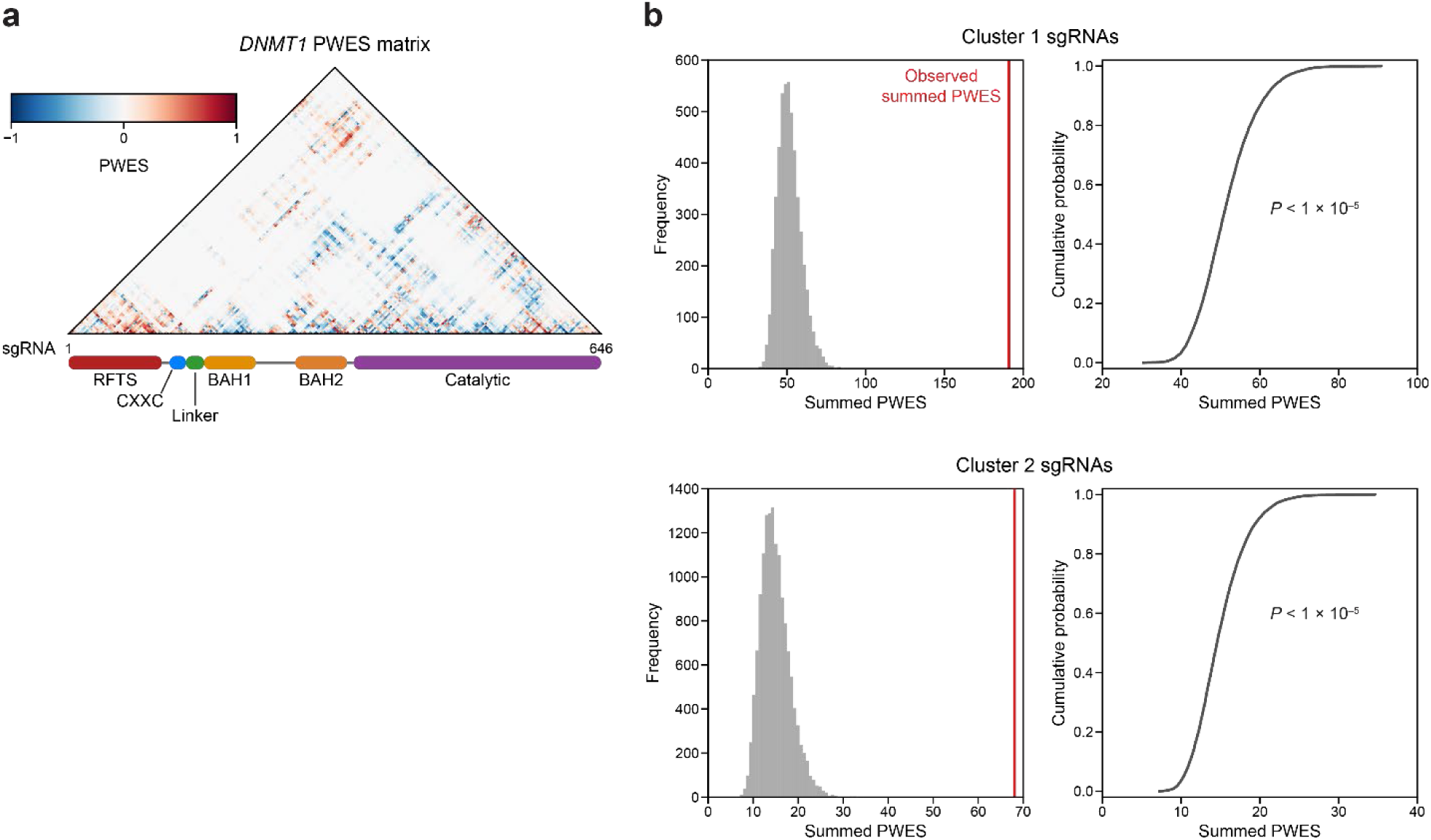
Linear and spatial clustering of CRISPR scanning data identifies putative hotspots in DNMT1 that modulate enzyme activity. **a**, Heatmap depicting the PWES matrix of all pairwise combinations of sgRNAs (*n* = 646) targeting resolved residues in the structure of autoinhibited DNMT1 (PDB: 4WXX). sgRNAs are ordered by their targeted amino acid position on the DNMT1 coding sequence. **b**, Histograms (left) and corresponding empirical cumulative distribution functions (ECDF, right) showing the simulated distribution of summed PWES values for cluster 1 (top) and cluster 2 (right) and the observed summed PWES value (red line). Summed PWES values were calculated as the sum of the absolute values of all intra-cluster PWES values. The simulated distribution was generated by shuffling the targeted amino acid positions of the sgRNAs and recalculating the summed PWES for *n* = 10,000 iterations. Empirical *P* values were calculated for cluster 1 and cluster 2 as 1 − ECDF(observed summed PWES value) and shown on the ECDF plots (right).

**Supplementary Figure 3.**
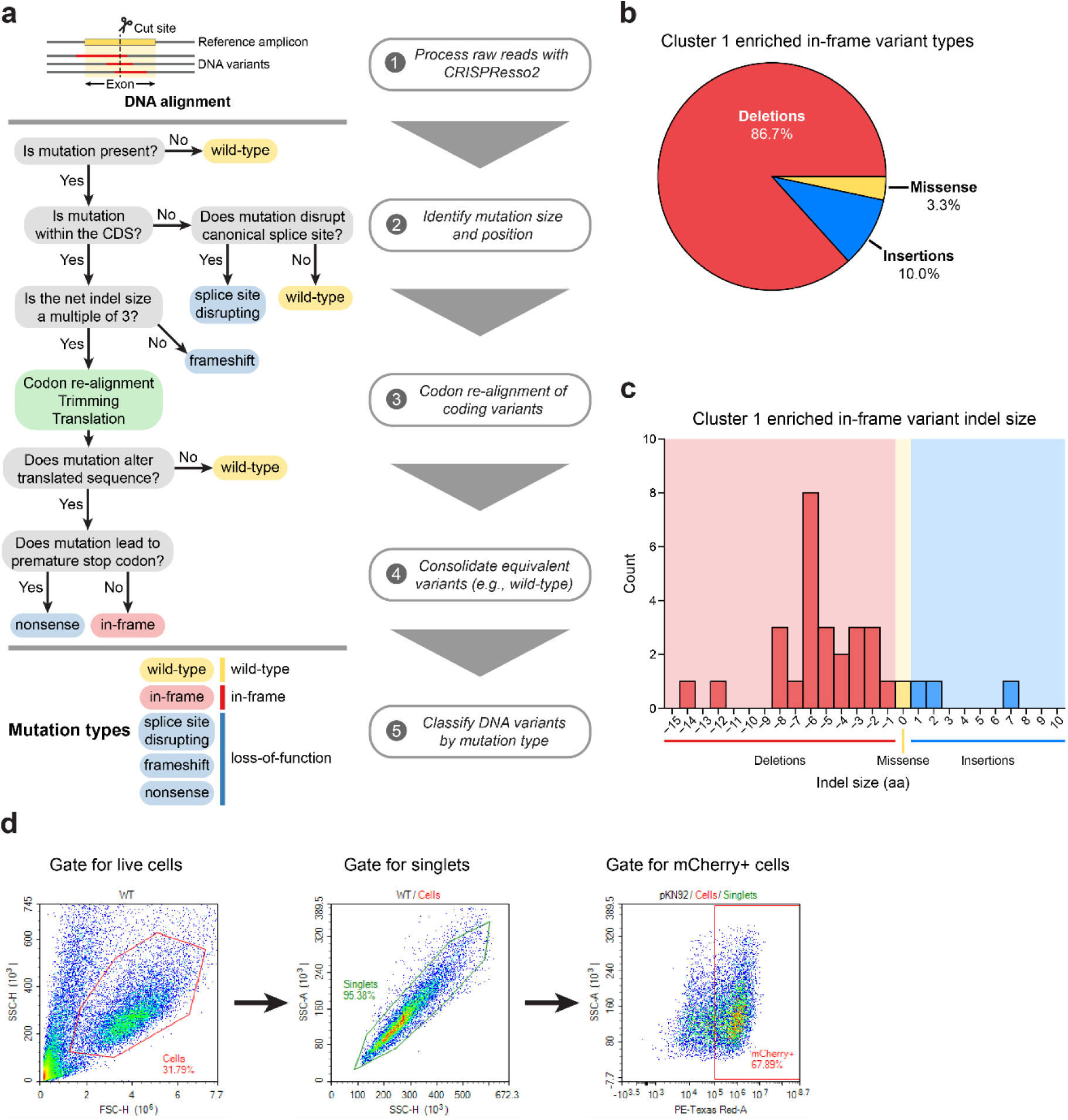
Cluster 1 sgRNAs generate hypermorphic DNMT1 mutations in the RFTS, CXXC, and linker regions that abrogate autoinhibition. **a**, Overview of the individual sgRNA mutational profiling and variant identification workflow. Raw sequencing data were first processed with CRISPResso2^35^ to align and quantify DNA variants. These DNA variants were then processed by a custom pipeline and classified by mutation type (represented as a decision tree in steps 2–4). In-frame variants were re-aligned, trimmed, and translated to generate the protein variant alignment tables shown in **Figure 3** and **Figure 5**. Splice site disrupting, frameshift, and nonsense mutations were combined into a single ‘loss-of-function’ mutation type for downstream analysis. **b**, Pie chart depicting the mutation type breakdown (e.g., insertion, deletion, missense) of enriched in-frame variants (*n* = 30) found in the selected cluster 1 sgRNAs (see **Fig. 3a,d**). In-frame variants were considered enriched if the observed frequency in DAC treatment was ≥1% and the log_2_(fold-change) of the observed frequency in DAC versus vehicle treatment was ≥2. sgT1503 and sgG1504/N1505 were excluded due to the lack of enriched in-frame variants. **c**, Bar chart showing the observed frequency (counts, *y*-axis) distribution of insertion (right, blue) and deletion (left, red) sizes (*x*-axis) for the enriched in-frame variants found in cluster 1 sgRNAs (from **b**). Missense mutations are highlighted in yellow. **d**, Flow cytometry pseudocolor density plots demonstrating the representative gating strategy used in the cellular fluorescent protein reporter assays to select mCherry-positive cells for analysis. The plots shown are taken from K562 cells expressing WT DNMT1–EGFP–IRES–mCherry after 3 d of vehicle treatment. Plots are representative of two independent experiments.

**Supplementary Figure 4.**
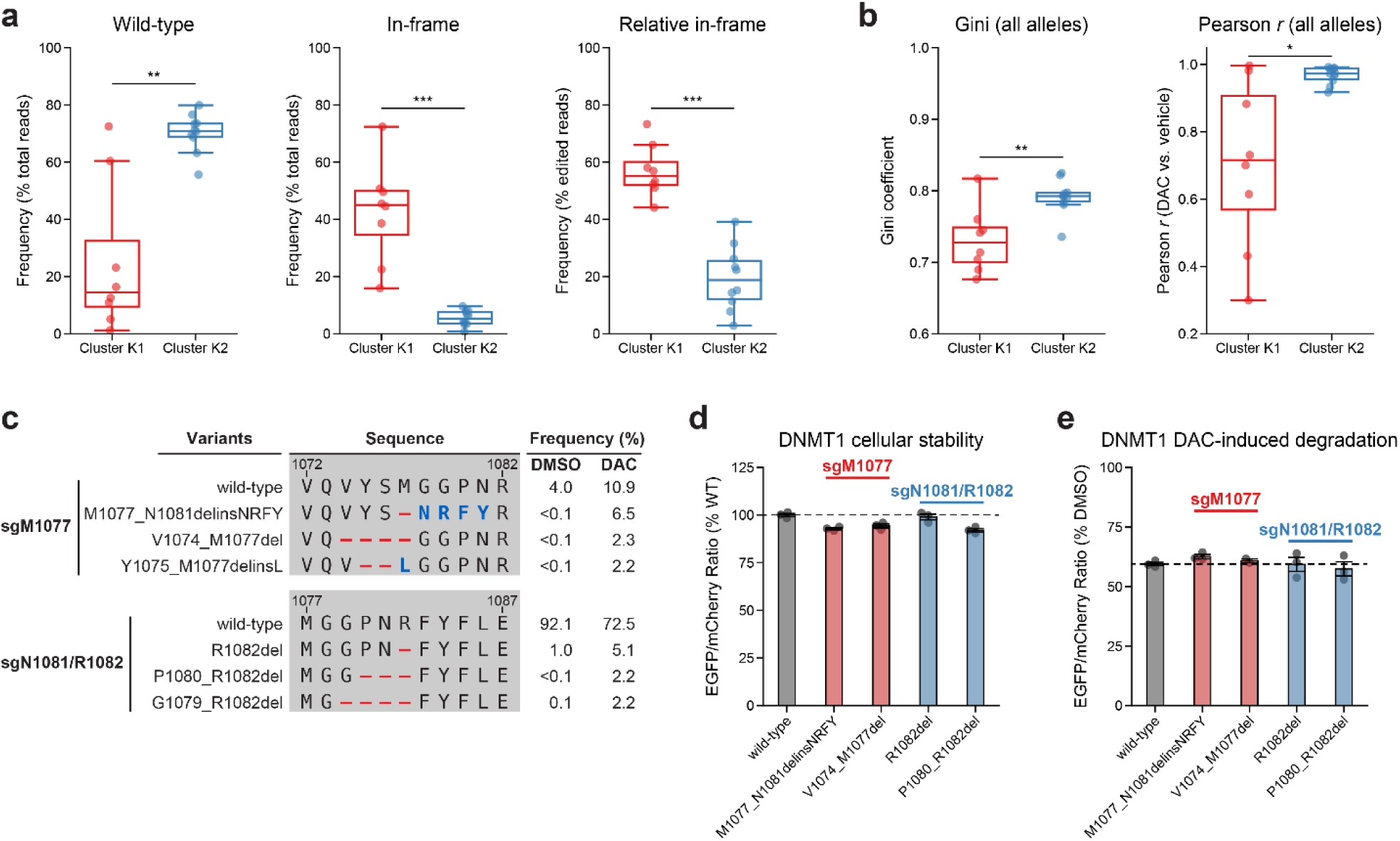
Integrative analysis reveals distinct mutational profiles between cluster 1 and cluster 2 sgRNAs. **a–b**, Box plots comparing various mutational profile metrics (*y*-axes) between cluster K1 (*n* = 8, red) and K2 (*n* = 10, blue) sgRNAs (*x*-axes). Individual sgRNAs are plotted as points. The central band, box boundaries, and whiskers represent the median, interquartile range (IQR), and 1.5 × IQR, respectively. *P* values (\**P* ≤0.05; \*\**P* ≤0.01, \*\*\**P* ≤0.001) were calculated with two-sided Mann-Whitney tests. **a**, The observed frequencies of wild-type (left) and in-frame (middle, right) variants in DAC treatment. Wild-type frequencies were calculated as the percentage of total reads. In-frame variant frequencies were calculated as a percentage of total reads (middle) and relative to the number of edited reads (right). **b**, The Gini coefficient of the observed allele frequency distribution in DAC treatment (left) and Pearson correlations of variant frequencies in DAC versus vehicle treatment (right). Gini coefficients and Pearson correlations were calculated using the observed frequencies (percentage of total reads) of all alleles. **c**, Table showing the amino acid sequence alignment and observed frequencies (percentage of total reads) of the wild-type and top enriched in-frame variants in the BAH2-targeting sgRNAs sgM1077 (top) and sgN1081/R1082 (bottom) after 8 weeks of treatment with vehicle (DMSO) or 100 nM decitabine (DAC). In-frame variants were considered enriched if the observed frequency in DAC treatment was ≥1% and the log_2_(fold-change) of the observed frequency in DAC versus vehicle treatment was ≥2. Enriched in-frame variants meeting these criteria were sorted by their observed frequency in DAC treatment and the top three most abundant are shown. Amino acid deletions are represented as red dashes and substitutions are highlighted in blue. **d**, Bar plot showing the cellular stability of wild-type DNMT1 and selected BAH2 variants (*x*-axis) enriched in sgM1077 (red) and sgN1081/R1082 (blue) in K562 cells as measured by the EGFP–mCherry fluorescence reporter system. Stability was calculated as the EGFP/mCherry ratio of the construct normalized to the EGFP/mCherry ratio of wild-type DNMT1 after 3 d of vehicle treatment (percentage of wild-type EGFP/mCherry ratio, *y*-axis). The dotted line represents the mean stability of the wild-type DNMT1 construct. Bars represent the mean ± s.d. across three replicates. One of two independent experiments is shown. **e**, Bar plot showing the degradation of wild-type DNMT1 and selected BAH2 variants (*x*-axis) enriched in sgM1077 (red) and sgN1081/R1082 (blue) in K562 cells as measured by the EGFP–mCherry fluorescence reporter system. Degradation was calculated for each construct as the EGFP/mCherry ratio of the construct in DAC treatment normalized to vehicle treatment (percentage of DMSO EGFP/mCherry ratio, *y*-axis). Cells were treated with vehicle or DAC for 3 d. The dotted line represents the mean degradation of the wild-type DNMT1 construct. Bars represent the mean ± s.d. across three replicates. One of two independent experiments is shown.

**Supplementary Figure 5.**
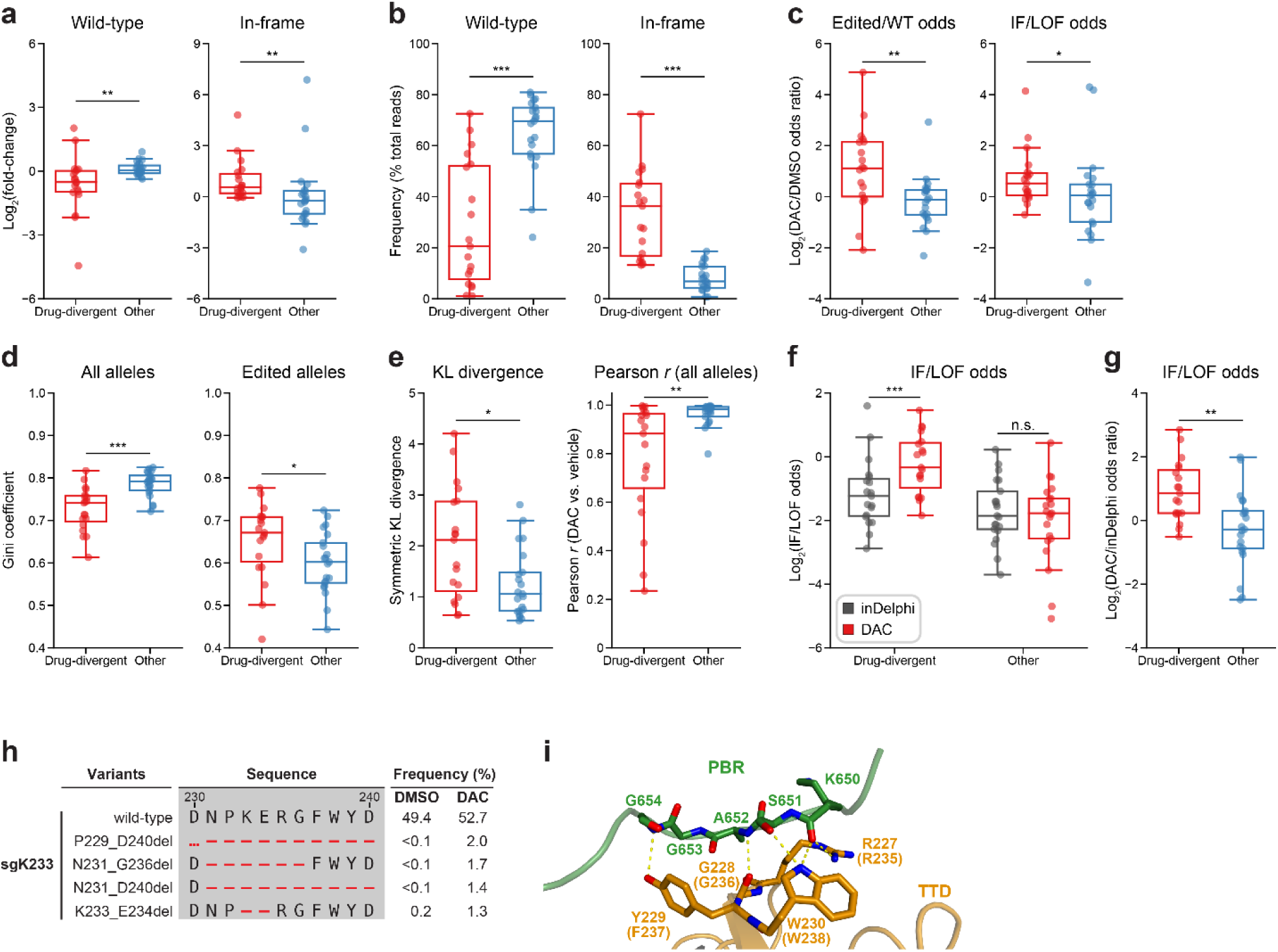
Mutational profiling analysis of individual sgRNAs nominates *UHRF1*-targeting sgRNAs with predicted gain-of-function outcomes. **a–e**, Box plots comparing various mutational profile metrics (*y*-axes) between ‘drug-divergent’ (*n* = 19, red) or ‘other’ (*n* = 21, blue) sgRNAs (*x*-axes). Individual sgRNAs are plotted as points. The central band, box boundaries, and whiskers represent the median, interquartile range (IQR), and 1.5 × IQR, respectively. *P* values (\**P* ≤0.05; \*\**P* ≤0.01, \*\*\**P* ≤0.001) were calculated with two-sided Mann-Whitney tests. **a**, The log_2_(fold-change) enrichment of wild-type (left) and in-frame variants (right) in DAC versus vehicle treatment. Log_2_(fold-change) was calculated based on the observed frequency (percentage of total reads) in DAC versus vehicle treatment. **b**, The observed frequencies (percentage of total reads) of wild-type (left) and in-frame (right) variants in DAC treatment. **c**, The log_2_(odds ratio) in DAC versus vehicle treatment for edited versus wild-type odds (left) and in-frame versus loss-of-function (LOF) odds (right). Edited/wild-type odds were calculated using absolute frequencies (percentage of total reads) and in-frame/LOF odds were calculated using relative frequencies (percentage of edited reads). **d**, The Gini coefficients of the observed allele frequency distribution in DAC treatment calculated using the observed frequencies of all alleles (percentage of total reads, left) or edited alleles (percentage of edited reads, right). **e**, The symmetric Kullback-Leibler (KL) divergence (left) and Pearson correlations (right) of observed allele frequencies in DAC versus vehicle treatment. The symmetric KL divergence and Pearson correlations were calculated with the observed frequencies (percentage of total reads) of all alleles. **f**, Box plots showing the log_2_(in-frame/LOF odds) (*y*-axis) for drug-divergent (red) and other (blue) sgRNAs (*x*-axis) as predicted by inDelphi (gray) and as observed in DAC treatment (red). Individual sgRNAs are plotted as points. The central band, box boundaries, and whiskers represent the median, interquartile range (IQR), and 1.5 × IQR, respectively. *P* values (\*\*\**P* ≤0.001; n.s., not significant) were calculated with two-sided Wilcoxon signed-rank tests. **g**, Box plots showing the log_2_(odds ratio) for in-frame/LOF odds in DAC treatment versus inDelphi predictions (*y*-axis) for drug-divergent (red) and other (blue) sgRNAs (*x*-axis). Individual sgRNAs are plotted as points. The central band, box boundaries, and whiskers represent the median, interquartile range (IQR), and 1.5 × IQR, respectively. The *P* value (\*\**P* ≤0.01) was calculated with a two-sided Mann-Whitney test. **h**, Table showing the amino acid sequence alignment and observed frequencies (percentage of total reads) of the wild-type and top enriched in-frame variants observed in sgK233 targeting the TTD domain of UHRF1 after 8 weeks of treatment with vehicle (DMSO) or 100 nM decitabine (DAC). In-frame variants were considered enriched if the observed frequency in DAC was ≥1% and the log_2_(fold-change) of the observed frequency in DAC versus vehicle treatment was ≥2. All enriched in-frame variants meeting these criteria are shown and ordered by their observed frequency in DAC treatment. Amino acid deletions are represented as red dashes. Red ellipses are used to denote amino acid deletions that exceed the length of the shown sequence alignment. **i**, Structural view of the zebrafish UHRF1 TTD domain (gold) complexed to a human UHRF1 PBR peptide (green) showing key residues in the TTD (human UHRF1 residues R227–W230) that form inter-domain contacts with the PBR and are targeted by enriched in-frame variants found in sgK233 (from **h**). Key residues forming polar contacts (yellow) are highlighted as sticks and annotated. For TTD residue annotations, the upper and lower (in parentheses) text indicate the residue identity and position in zebrafish and human UHRF1, respectively. (PDB: 6B9M)

